# *Pseudomonas putida* JM37 as a novel bacterial chassis for ethylene glycol upcycling

**DOI:** 10.1101/2025.10.07.680640

**Authors:** Francisco J. Molpeceres-García, Alejandro García-Miró, Elvira Mateos, Alicia Prieto, David Sanz, José I. Jiménez, Jorge Barriuso

**Affiliations:** Department of Biotechnology, Center for Biological Research Margarita Salas, Spanish National Research Council (CIB-CSIC), 28040 Madrid, Spain; Department of Life Sciences, Imperial College London, South Kensington Campus, London SW7 2AZ, United Kingdom

**Keywords:** *Pseudomonas putida*, ethylene glycol metabolism, upcycling, synthetic biology chassis

## Abstract

Ethylene glycol (EG), one of the main monomers of polyethylene terephthalate (PET), is an attractive target for microbial upcycling. Despite this interest, there is a limited number of described organisms that can efficiently metabolise EG. Here, we report the metabolic and biotechnological potential of *Pseudomonas putida* JM37 as a novel bacterial chassis for EG valorization. We show that JM37 efficiently grows on EG as the sole carbon and energy source, outperforming other *Pseudomonas* strains. Genome sequencing and directed mutagenesis revealed that genetic redundancies in the glyoxylate assimilation pathways underlie its robust EG metabolism. Beyond biomass generation, we demonstrated the biotechnological potential of JM37. This strain was able to accumulate medium-chain polyhydroxyalkanoates (mcl-PHAs), dominated by C10 monomers, directly from EG. Moreover, JM37 successfully expressed heterologous biosynthetic pathways, including a violacein biosynthetic operon and a PET-hydrolase which has been secreted actively into the extracellular medium. Together, our results support the use of *P. putida* JM37 as a versatile synthetic biology chassis for sustainable EG upcycling and as a promising platform for circular bioproduction.

## Introduction

Ethylene glycol (EG) is a C_2_ diol extensively employed in several industrial applications as antifreezing, aircraft and runway de-icing, engine coolants, natural gas processing and polyester synthesis (Shehata et al., 2024). Globally, most of the EG is destined to the fabrication of the thermoplastic polyethylene terephthalate (PET) whose production is estimated to reach 61 million tons per year by 2060 (OECD, 2022; Pang et al., 2016). Most of PET products are single-use, as this plastic is mainly employed in packaging industry, and its recycling rates are as low as ∼21% in the US, mainly due to the lower quality and higher cost of recycled PET compared to virgin PET (Kim et al., 2019a). Recently, biotechnological approaches to plastic waste management have focused on PET monomers, terephthalic acid (TPA) and EG, as substrates for biological upcycling, i.e., their conversion to value-added products, such as flavourings like vanillin (Sadler & Wallace, 2021) or vanillic acid (Kim et al., 2019b); chemicals with pharmacological applications like lycopene (Diao et al., 2023); painkillers like paracetamol (Johnson et al., 2025); industrial chemicals like glycolic acid (Carniel et al., 2023) or bioplastics like polyhydroxyalkanoates (PHA) (Kenny et al., 2008; Tiso et al., 2021).

There are two main approaches to obtain TPA and EG from PET: chemical catalysis and biological catalysis. In the chemical catalysis approach, PET is depolymerized to its fundamental monomers or to intermediates like dimethyl terephthalate or bis(2-Hydroxyethyl) terephthalate (BHET), employing alcohols (alcoholysis), amines (aminolysis), strong acids or bases (hydrolysis) or EG (glycolysis). Although this process is very quick and efficient, it requires high temperatures and, frequently, toxic catalysts (Carniel et al., 2024). The biological catalysis approach consists on an enzymatic depolymerization employing PET-hydrolases expressed by microorganisms. These enzymes consist mainly in lipases, cutinases, esterases or carboxylesterases (Kawai et al., 2024). This approach does not need of strong solvents, toxic catalysts or high energy consumption, resulting in a more environmentally friendly process. Nevertheless, reaction rates are much slower, and properties of the polymer such as crystallinity suppose a barrier for the depolymerization, because enzymes only attack efficiently amorphous regions of the plastic (Carniel et al., 2024).

Given the interest in PET monomers upcycling and the widespread use of EG, the study of microorganisms able to metabolize this diol is of great interest. *Mycobacterium* sp. E44 and *Acetobacterium woodii* are able to grow using EG (Trifunović et al., 2016; Wiegant & De Bont, 1980). In aerobic bacteria, the most studied pathway involves the oxidation of EG to glycolaldehyde, then to glycolate and finally to glyoxylate (Boronat et al., 1983; Hachisuka et al., 2022; Ren et al., 2025; Shimizu et al., 2024). Glyoxylate can enter the central metabolism through the glyoxylate shunt or dedicated pathways, allowing the generation of biomass and/or energy (Ren et al., 2025).

The model organism *Pseudomonas putida* KT2440 is able to assimilate EG via this oxidative pathway (Fig. 1). In this strain, the pyrroloquinoline quinone (PQQ)-dependent periplasmic alcohol dehydrogenases PedE and PedH can be used for the initial oxidation of EG to glycolaldehyde. In a next step, two aldehyde dehydrogenases (PedI and AldB) convert glycolaldehyde into glycolate, which is oxidised to glyoxylate by the membrane anchored glycolate oxidase GlcDEF. Glyoxylate can be metabolised through the glyoxylate shunt, either by condensation with an acetyl-CoA to form malate by the malate synthase GlcB, or with succinate to form D-isocitrate by the isocitrate lyase AceA. Alternatively, glyoxylate can be transformed to glycerate and then to pyruvate through the glyoxylate carboligase pathway. This pathway involves a glyoxylate carboligase (Gcl), a hydroxypyruvate isomerase (Hyi), a tartronate semialdehyde reductase (GlxR), a hydroxypyruvate reductase (TtuD), a glycerate kinase (GarK) and a pyruvate kinase (TtuE) (Franden et al., 2018). These genes are also present in *Pseudomonas umsongensis* GO16, which is able to grow on EG (Narancic et al., 2021), suggesting their conservation among the *Pseudomonas* genus. Although all this metabolic machinery is present in *P. putida* KT2440, this strain needs genetic modifications to generate biomass from EG (Franden et al., 2018; Li et al., 2019). Moreover, *P. putida* KT2440 has been recently engineered to express the β-hydroxyaspartate cycle from *Paracoccus denitrificans*, which allows to generate biomass by the condensation of two glyoxylates into oxaloacetate employing less NADH and without carbon loss in form of CO_2_ (Schada von Borzyskowski et al., 2023).

**Figure 1.**
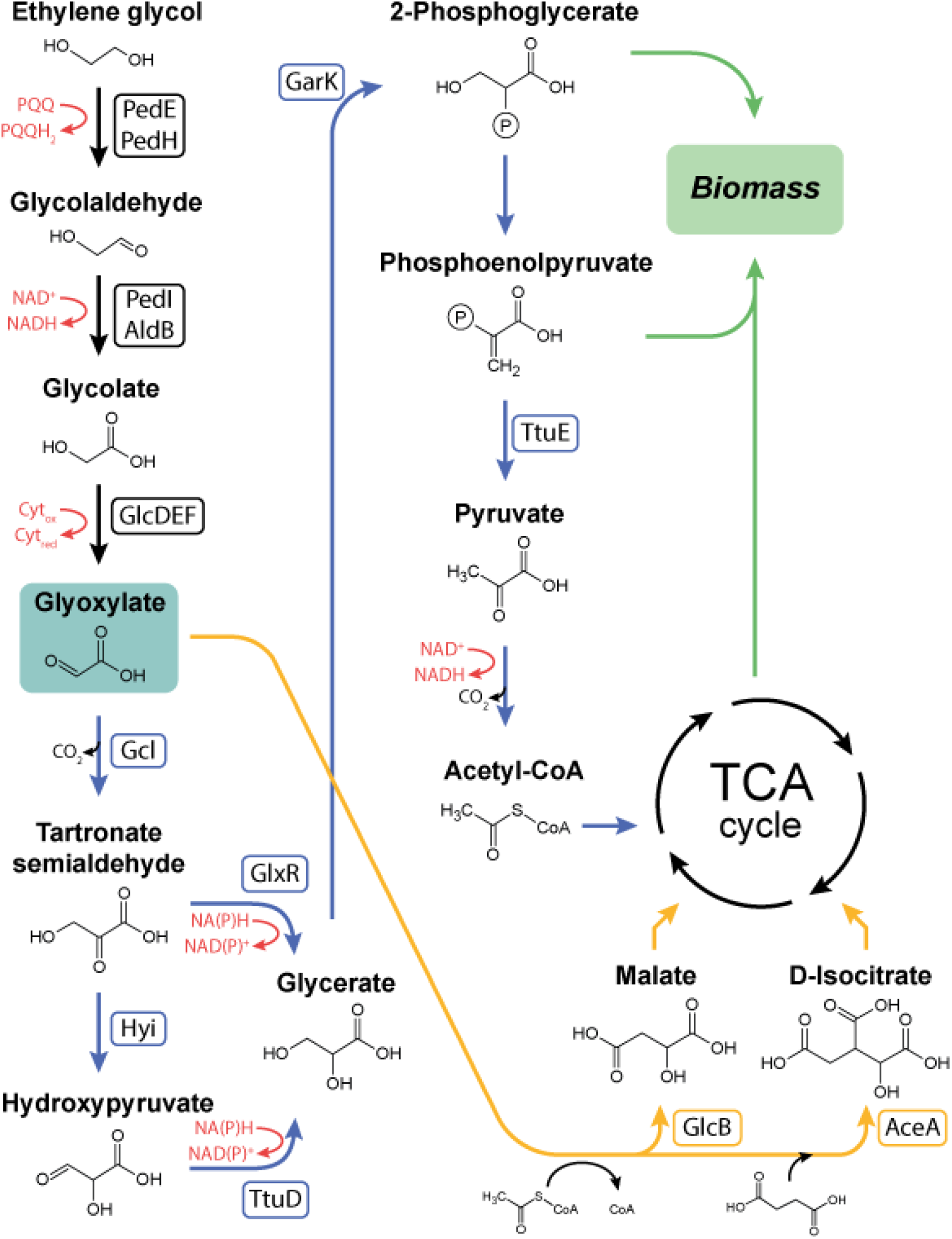
Ethylene glycol metabolism scheme in *P. putida*. Blue represents the Gcl pathway. Yellow represents the glyoxylate shunt. Red represents redox reactions.

The strain *P. putida* JM37 can efficiently grow on EG as the only carbon an energy source (Mückschel et al., 2012). However, it is not clear why this strain behaves differently to KT2440. In this work, we have sequenced and studied the genome of *P. putida* JM37 and found that contains an operon formed by *glcDEF* and the Gcl pathway, as well as a duplication in *glcB*. To elucidate whether JM37 assimilates EG through Gcl and the glyoxylate shunt, we disrupted the key genes in these pathways employing a CRISPR-Cas9 base editing approach (Volke et al., 2022). Furthermore, we investigated the capacities of this strain for the upcycling of PET-derived EG. We demonstrated JM37’s capacity for growing at high concentrations of EG and accumulating medium-chain polyhydroxyalkanoates (mcl-PHA). In addition, we were able to express in this bacterium a violacein biosynthesis operon and a secreted PET-hydrolase, demonstrating the capacity of this strain to produce heterologous value-added compounds and express heterologous active enzymes. Our work delves into *P. putida* JM37 EG metabolism and postulates this strain as a synthetic biology chassis.

## Materials and methods

### Microorganisms and culture conditions

The bacterial strains and plasmids used in this work are listed in Table 1. *P. putida* JM 37 strain was kindly provided by Dr. Bernhard Hauer (University of Stuttgart, Germany). Unless indicated, microorganisms were grown in LB medium and transferred to LB or MC minimal medium: KH_2_PO_4_ 0.33 g/L, Na_2_HPO_4_ 1.2 g/L, NH_4_Cl 0.1 g/L, MgSO_4_*7H_2_O 0.1 g/L, CaCl_2_ 44 mg/L, adjusted to pH 7. The vitamins and trace elements solutions employed are described in (Barragán et al., 2004). Three biological replicates were used in all experiments. When necessary, 50 µg/mL of kanamycin and 35 µg/mL of gentamicin were added to the growth media. Cultures were grown at 30 ºC and 200 rpm and gene expression was induced with 1 mM IPTG.

**Table 1.**
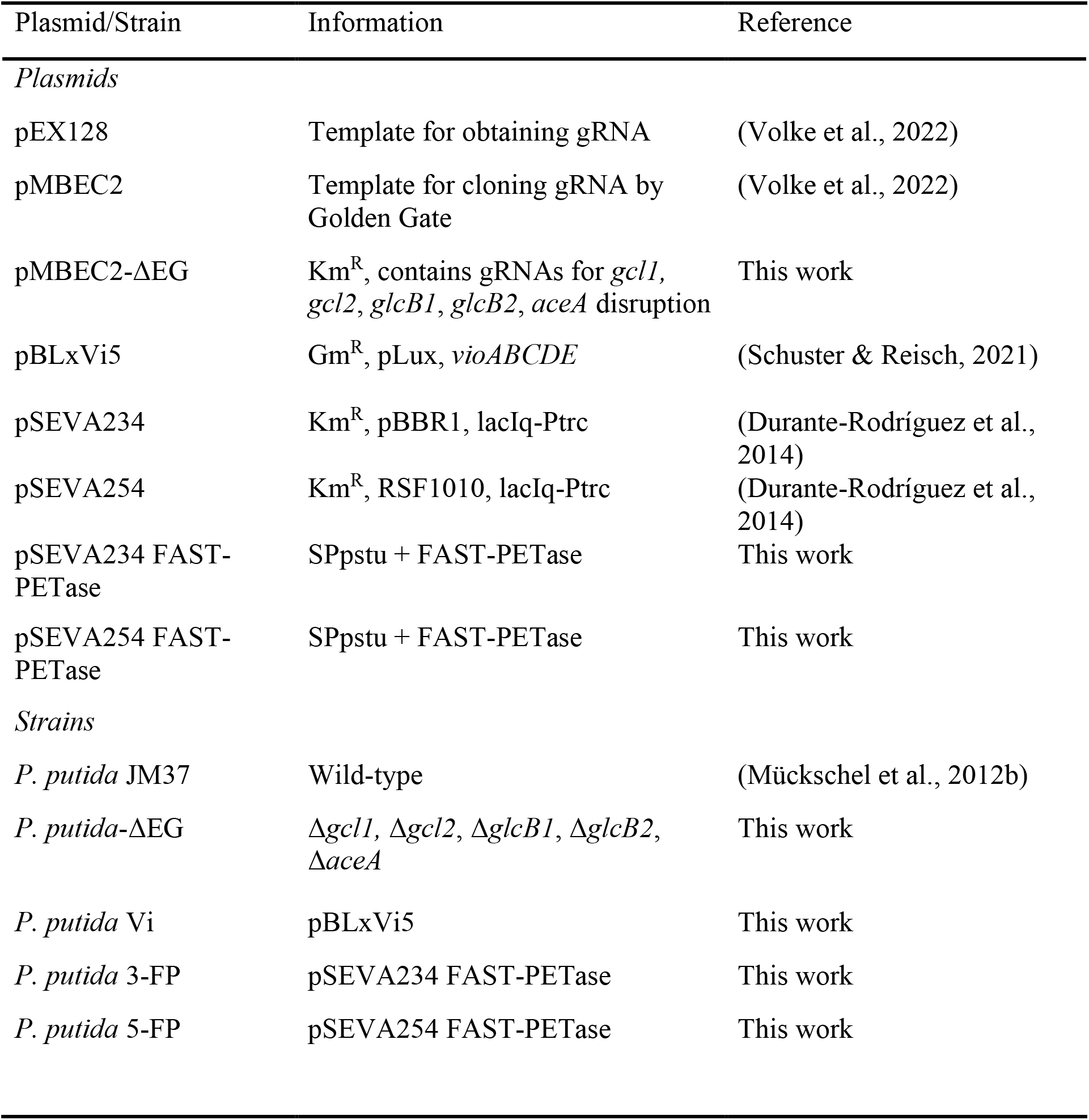
Strains and plasmids employed in this study.

### Growth of P. putida JM37in EG

*P. putida* JM37 was grown in MC minimal medium with EG (Sigma-Aldrich, REF 102466) as sole carbon source in concentrations ranging from 10 mM to 2 M. Growth curves were performed in a 96 well plate employing a Victor Nivo Multimode Plate Reader (PerkinElmer) and in 250 mL rotatory flasks filled with 50 mL of media, in both cases at 30 ºC and 200 rpm. To study the influence of nitrogen in growth, we employed 0.3 g/L of NH4CL with the higher EG concentrations (200 mM, 500 mM and 1 M). Growth was measured as optical density at 600 nm (OD^600^).

### Genome sequencing and analysis

*P. putida* JM37 genome was sequenced by MicrobesNG (Birmingham, UK) using an Illumina NovaSeq 6000 platform (coverage ∼100X). Raw reads (FASTQ) were deposited at NCBI SRA repository under accession number PRJNA1330276. The assembly was performed by the company, obtaining 154 contigs. Annotation was done at RAST server (Aziz et al., 2008) and EG degradation genes were found using *P. putida* KT2440 as reference. *P. putida* JM37 genes were compared with those of *P. putida* KT2440 and *P. umsongensis* GO16 employing RAST inner BLASTp.

### CRISPR-Cas9 base editing editing

To confirm the key genes used by *P. putida* JM37 to metabolise EG, we selective knocked out the two copies of the glyoxylate carboligase (*gcl*), the two copies of the malate synthase (*glcB*) and the only copy of the isocitrate lyase (*aceA*). These mutations were performed by a CRISPR-Cas9 assisted method developed by Volke et al. (2022). Briefly, this technique allows C**→**T substitutions, generating premature STOP codons and thus interrupting the translation of the Open Reading Frame. CRISPy-web service (Blin et al., 2016) was used to obtain gRNA sequences for our target mutations, which were used to design primers. With the gRNA-primers, PCRs were performed with the pEX128 plasmid as template, obtaining gRNAs with a Golden Gate consensus sequence. These were cloned into the pMBEC2 plasmid (kanamycin resistance) (Volke et al., 2022) following authors’ protocol. As a result, the plasmid pMBEC2-ΔEG, which contained gRNAs for the five knock-outs, was obtained. This plasmid was transformed into *Escherichia coli* DH10β by heat shock and plated in LB with kanamycin 50 µg/mL for colony selection. Positive colonies lacked GFP expression. To confirm the correct integration of the gRNAs, plasmid was extracted with a Monarch® Plasmid Miniprep Kit (New England Biolabs) and sequenced by Full Circle Labs (London, UK). *P. putida* JM37 was transformed by electroporation with the pMBEC2-ΔEG plasmid, resulting in the strain *P. putida* JM37-ΔEG.

To confirm the mutations, *P. putida* JM37-ΔEG was grown in LB with kanamycin and inoculated at 0.1 OD^600^ in MC medium with EG 10 mM, then grown for 48 h at 30 ºC and 200 rpm. The wild type strain was used as positive control. Furthermore, the genome of *P. putida* JM37-ΔEG was sequenced by MacroGen (Seoul) with Illumina NovaSeq X. Raw data was provided as FASTQ (Paired-end). For FASTQ processing, fastp (Chen et al., 2018) was employed to trim polyG and adapter sequences, as well as to eliminate < 30 Phred quality score and < 30 bp reads. The genome was assembled in contigs using SPAdes (Prjibelski et al., 2020) (k-mers 21, 33, 55, 77). To check the genetic editions, native and mutated genes were aligned with Clustal-Omega (Madeira et al., 2024). CRISPR primers and aligned sequences can be found in Table S1 and Fig. S3. Raw reads (FASTQ) were deposited at NCBI SRA repository under the accession number PRJNA1330276.

### Electroporation protocol

Electroporation was employed for the transformation of *P. putida* JM37. First, electrocompetent cells were prepared. *P. putida* JM37 was grown O/N in LB, and then inoculated in 50 mL of fresh LB at OD^600^ 0.1. After 2 - 2.5 h, when the OD^600^ value reached ∼ 0.8, cells were washed twice with 50 mL of chilled Milli-Q water with 10% glycerol. Finally, the centrifuged pellet was resuspended in 1 mL of 10% glycerol and divided in 100 µL aliquots, which were immediately stored at -80 ºC. Electroporation was performed in 0.2 cm cuvettes with a MicroPulse elctroporator (BioRad) (2.5 kV, 200 Ω, 25 µF) adding 100 ng of plasmid. Cells were recovered in LB for 2 h and plated in LB with kanamycin 50 µg/mL for colonies selection.

### PHA analysis and quantification

To study PHA accumulation, *P. putida* JM37 was grown in MC medium with EG 50 mM as sole carbon source. After 24 h, PHA production was confirmed visually by Nile Red staining (Fig. 4) (Zuriani et al., 2013). Briefly, 500 µL of culture were centrifuged at 7.000 rpm, resulting pellet was resuspended in 50 µL of PBS and Nile Red was added to a final concentration of 5 µg/mL. PHA granules were observed with a DM4 B microscope (Leica) equipped with a *p*E-300 Lite LED illuminator (CoolLED) and a DFC345 FX camera (Leica).

To analyse the composition and quantify total intracellular PHA, 2-5 mg of freeze-dried biomass were methanolized. For each biological sample, two technical replicates were included. Briefly, freeze-dried pellets were resuspended in 2 mL of methanol containing 15% of sulfuric acid. Then, 2 mL of 3-methylbenzoic acid, dissolved in chloroform at 0.5 mg/mL, were added as an internal standard. Samples were incubated at 100 ºC for 5 h. Reactions were stopped by cooling in ice. Samples were washed twice by adding 2 mL of Milli-Q water, centrifuging for phase separation and discarding the aqueous phase and the cellular debris. A third washing step was done with a saturated solution of Na_2_CO_3_ to eliminate sulfuric acid remains. Finally, a small amount of Na_2_SO_4_ was added to completely remove water content.

Methanolized samples were analysed by GC-MS with an Agilent series 7890A gas chromatograph coupled with a 5975C MS detector (EI, 70 eV) and a DB-5HTDB-5HT column (400 ºC: 30 m x 0.25 mm x 0.1 µm film thickness). Retention times for the different methyl ester monomers were: 3.5 min (C6), 5.2 min (C8), 7.0 min (C10), 9.1 min (C12:1) and 9.4 min (C12). Internal standard was detected at 4.9 min.

### EG quantification by HPLC

An Agilent 1200 series HPLC was used to measure EG. Samples were centrifuged at 13.000 rpm and filtered with a 0.22 µm PTFE filter (Agilent) before analysis. EG was measured with an RID detector (Agilent) and a BioRad Aminex HPX-87H column (300 x 7.8 mm), employing H_2_SO_4_ 2 mM as mobile phase in an isocratic 0.5 mL/min flux for 25 min. The column was maintained at 60 ºC and RID at 35 ºC. EG was eluted at 19.5 min.

### PET-Hydrolase and Violacein Operon Cloning

Violacein synthesis operon *vioABCDE* was cloned using the plasmid pBLxVi5 (Gm resistance) (Schuster & Reisch, 2021), obtaining the *P. putida* Vi strain. The codon-optimized FAST-PETase (Lu et al., 2022) containing a SPpstu signal peptide from *Pseudomonas stutzeri* MO-19 (Fujita et al., 1989) was cloned into pSEVA234 (pBBR1) and a pSEVA254 (RSF1010) plasmids (Km resistance), using *Eco*RI and *Hind*III restriction sites (Durante-Rodríguez et al., 2014). Transformations were conducted by electroporation. The resulting strains were named *P. putida* 3-FP and *P. putida* 5-FP (Table 1).

### Violacein production and quantification

For crude violacein (violacein + deoxyviolacein) production, *P. putida* Vi was cultivated in MC medium with EG 50 mM as carbon source. For induction, 10 µM of N-(β-Ketocaproyl)-L-homoserine lactone (Sigma-Aldrich, REF K3007) was used. *P*. putida Vi was streaked in an MC-agar plate to check crude violacein production visually. For quantification, crude violacein was extracted and quantified following Schuster & Reisch (2021) protocol, with some modifications, after 18 h of cultivation in shake flasks. Briefly, 1 mL of culture was centrifuged at 12 000 × g for 5 min. For violacein extraction, the pellet was resuspended in 1mL of absolute ethanol containing 1% (v/v) acetic acid and incubated at 60 ºC and 1000 rpm for 10 min. For quantification, the solution was pelleted and the supernatant was measured at 570 nm in a UV 1900i spectrophotometer (Shimadzu). The crude violacein yield was quantified employing violacein molar extinction coefficient (10.955 L/(g cm)) (Wang et al., 2009).

### PETase activity assay

To asses FAST-PETase activity, *P. putida* 3-FP and 5-FP strains were cultured in LB for an O/N and then inoculated at 0.1 OD^600^ in fresh LB with IPTG 1 mM. Kanamycin was used for plasmid maintenance. Both strains without IPTG and the wild-type were used as negative controls. After 48 h, the activity was measured with a *p*-nitrophenyl-butyrate (*p*NPB) colorimetric assay. The hydrolyzation of 1.5 mM of *p*NPB (Sigma-Aldrich, REF N9876) was followed by measuring the absorbance increase at 410 nm. The reaction was conducted in a 50 mM phosphate buffer pH 7.5 employing 100 µL of the cultures’ supernatants. One unit of activity (1 U) is defined as the amount of enzyme used to release 1 μmol of *p*-nitrophenol (ε410 =15,200 M^-1^ cm^-1^) per minute. The activity results were normalized by cultures’ OD^600^ values for a fair comparison.

## Results and discussion

### Utilization of Ethylene Glycol as the Sole Carbon Source by Pseudomonas putida JM37

*P. putida* JM37 was confirmed to efficiently grow employing EG as the sole carbon source, as originally reported (Mückschel et al., 2012). The bacterium achieved a maximum growth of 1.13 OD^600^ in 24 h with 50 mM EG in a 96-well plate. Greater concentrations showed a decrease in growth, and 2 M resulted inhibitory (Fig. 2A). Microplate cultures are known to be affected by oxygen limitations; hence we also grew the bacterium in shake flasks, achieving in this case 2.2 OD^600^ with 50 mM EG in 24 h. Higher carbon availability did not increase the maximum biomass under these conditions suggesting that existence of additional nutrient limitations. These were overcome by adding three times more nitrogen. With this modification, *P. putida* JM37 achieved 4.9 OD^600^ in 48 h with 200 mM EG, consuming 126 mM of the substrate in 48h (Fig. 2B and C).

**Figure 2.**
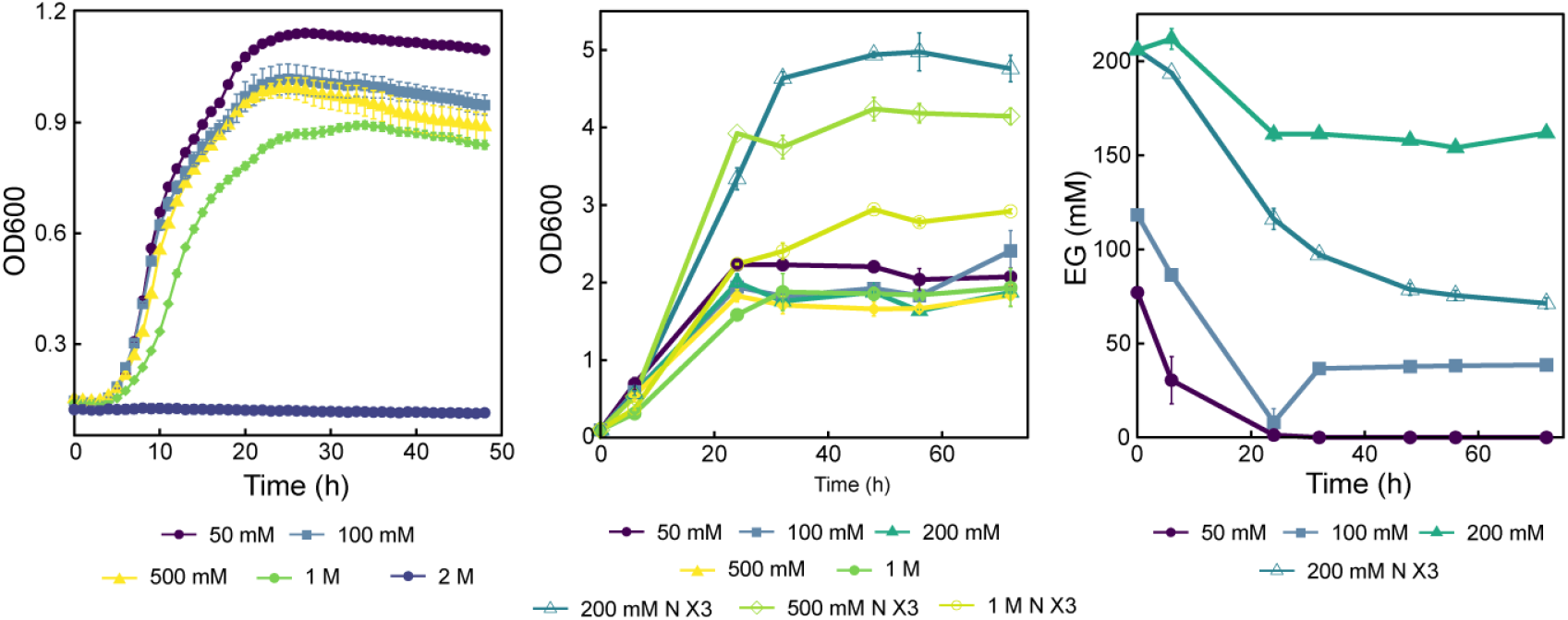
(A, B) Growth of *P. putida* JM37 in MC minimal medium with different concentrations of EG as the sole carbon source in (A) microplate reader with agitation and (B) shake flasks. (C) shows the consumption of different EG concentrations when growing in shake flasks. Results are given as the mean of n = 3 for (A) and n = 2 for (C). Error bars represent the standard error of these measurements.

Previous works have shown that *Acetobacterium woodi* DSM 1030 achieved 1.1 OD^600^ with 75 mM EG in 55 h in shake flask cultures (Trifunović et al., 2016). *Paracoccus denitrificans* can grow up to ∼1.1 OD^600^ in 48 h with 400 mM EG employing NAD-dependent oxidative enzymes (Ren et al., 2025). *E. coli* cannot naturally grow on EG, although an evolved strain derived from K12 with a mutated 1,2-propanediol oxidoreductase (FucO) is able to assimilate the substrate (Caballero et al., 1983).

The model organism *P. putida* KT2440 needs adaptative laboratory evolution (ALE) or genetic modifications to grow on EG. Li et al. (2019) demonstrated that the transcriptional regulator GclR is a repressor of the Gcl pathway in this strain. By either its deletion or a mutation in its binding site upstream *gcl*, KT2440 is able to grow on EG, highlighting the importance of this pathway for biomass generation from EG in this strain. Employing ALE, the authors obtained a mutant strain for GclR, PP_2046 and PP_2662 genes, encoding a LysR-type transcriptional regulator and a putative porin, respectively. This version of *P*. putida KT2440 could grow up to 0.6 cell dry weight (CDW) (g/L) in 24h with 26.7 mM EG in shake flasks.

With a different approach, Franden et al. (2018) designed a strain overexpressing the glycolate oxidase GlcDEF and the Gcl pathway. This modified *P. putida* KT2440 could tolerate up to 2 M EG and achieved a growth of almost 10 g/L DCW in 120 h, demonstrating the importance of GlcDEF in addition to the Gcl pathway for generating biomass in this strain.

Without any genetic modification, *P. umsongensis* GO16 is able to grow on EG as the sole carbon source. This strain achieved 0.4 g/L CDW in 48 h with 1.6 g/L (∼26 mM) EG (Narancic et al., 2021). After subdued to ALE, the strain *P. umsongensis* GO16 KS3 grew up to 3 OD^600^ in 10 h with 1.7 g/L (∼27 mM) EG as the sole carbon source in shake flasks (Tiso et al., 2021).

The growth on EG achieved by *P. putida* JM37 in our experiments is among the highest reported in literature, demonstrating the potential of this strain for developing EG upcycling strategies.

### Genetic Redundance of Pseudomonas putida JM37 Enable Efficient Ethylene Glycol Utilization

We found the homologous genes for EG assimilation from *P. putida* KT2440 and *P. umsongensis* GO16 in the genome of *P. putida* JM37, presenting a similar distribution in the chromosome: an operon containing *pedE, pedH, pedI*, and *aldB*; an operon with *glcDEF*, and an operon with the Gcl pathway (*gcl, hyi, glxR, ttuD, ttuE*). The genes *aceA* and *glcB* were not clustered, but they are present in different parts of the genome (Fig. 3). The operon containing *pedE, pedH, pedI*, and *aldB* is controlled by the transcriptional regulator *pedR1* and the PAS/PAC sensor hybrid histidine kinase *pedS1* in *P. putida* KT2440. In this case, both *pedR1* and *pedS1* were found in the genome of *P. putida* JM37, suggesting a similar regulation.

**Figure 3.**
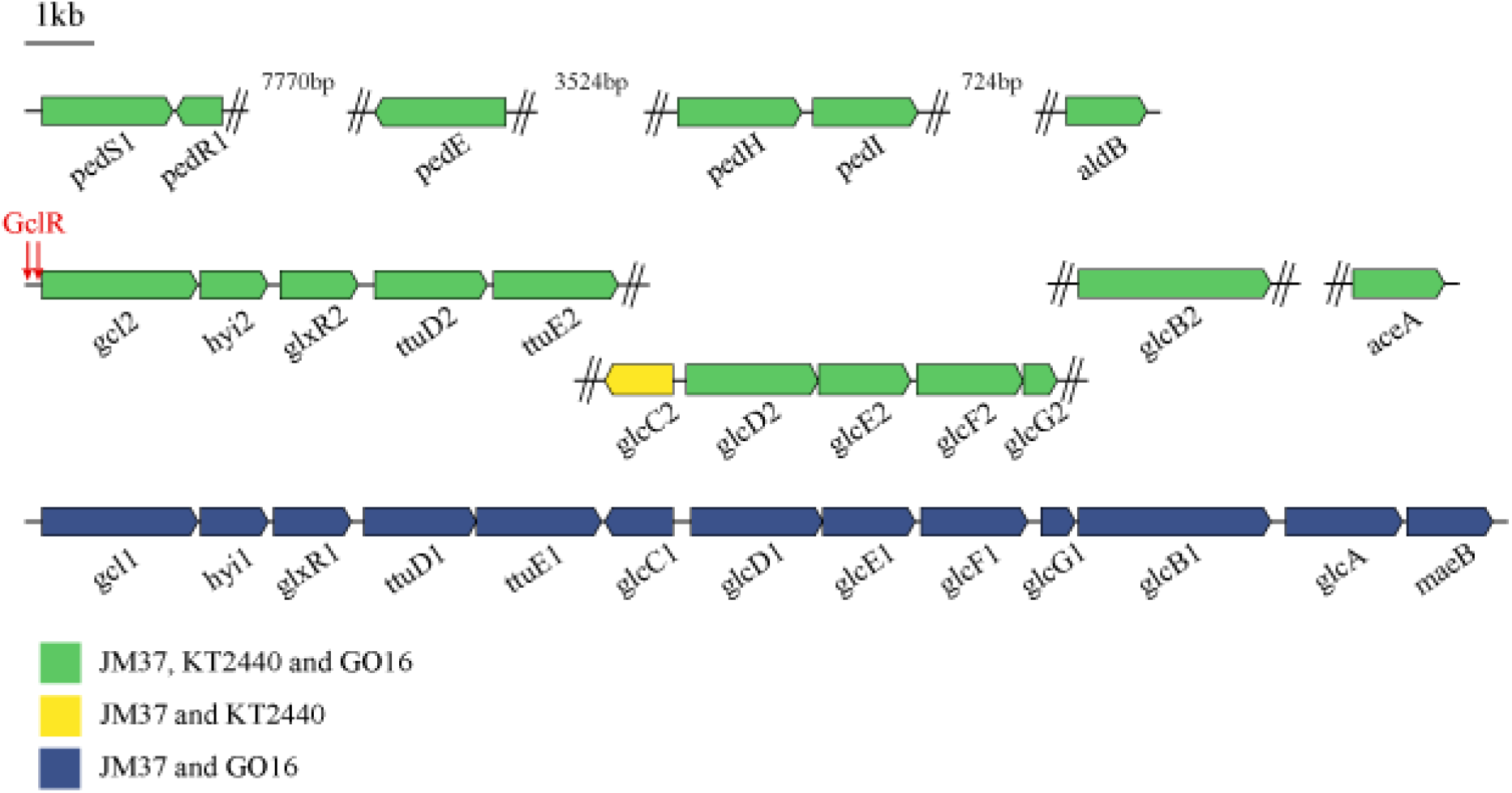
Disposition of genes related to EG metabolism in *P. putida* JM37. Common dispositions with *P. putida* KT2440 and *P. umsongensis* GO16 are explained in the legend. GclR putative binding sites are pointed in red. Gene sizes and distances are represented for *P. putida* JM37. Protein homologies are shown in Table 2.

Franden et al. (2018) demonstrated that the Gcl pathway, which is repressed by the regulator GclR (Li et al., 2019), is fundamental for biomass generation from EG in *P. putida* KT2440. Furthermore, the authors showed that *P. putida* KT2440 accumulates the toxic intermediate glycolaldehyde when metabolizing EG, and solved this problem by overexpressing GlcDEF, which dramatically increase the growth of *P. putida* KT2440 on this substrate.

**Table 2.**
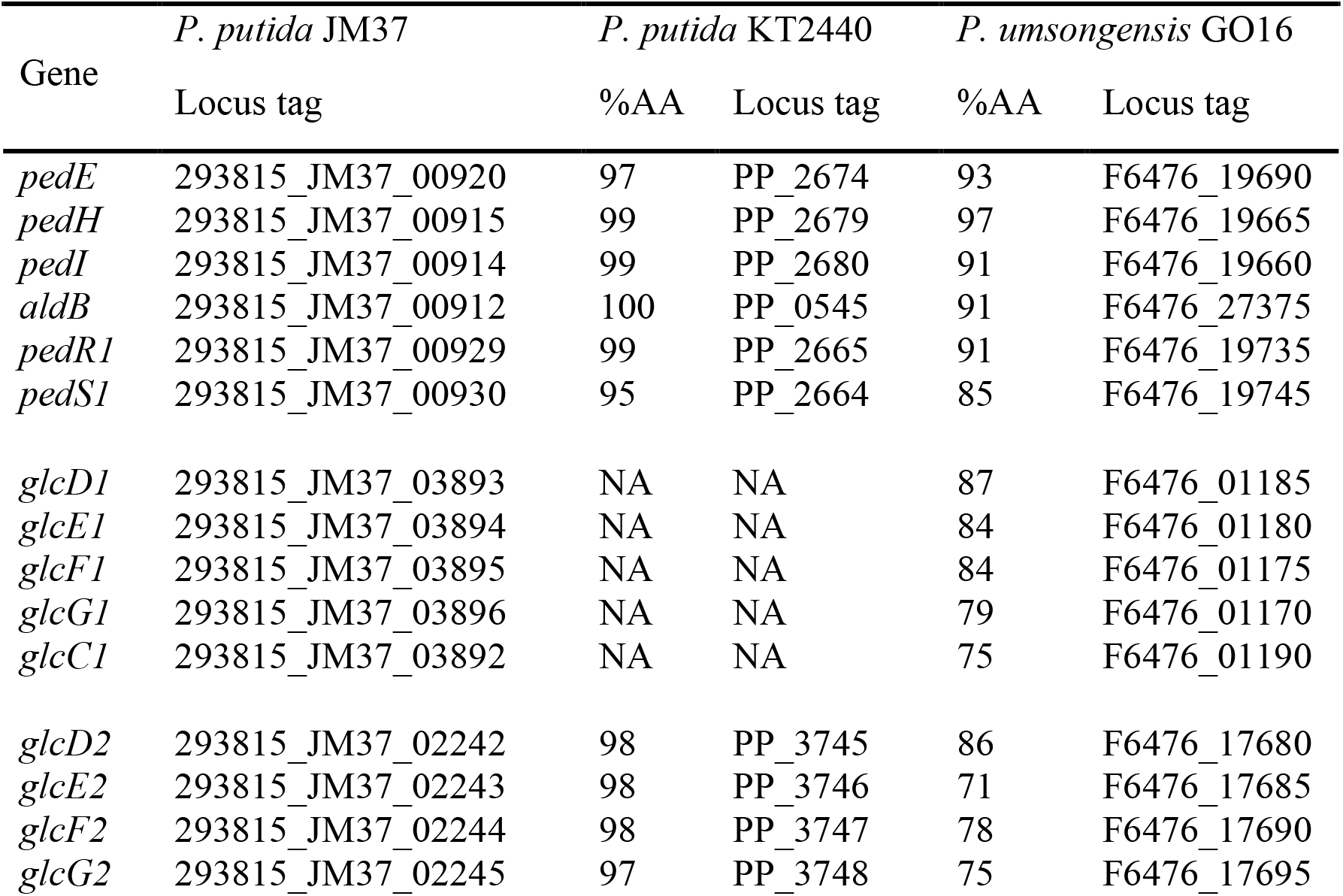
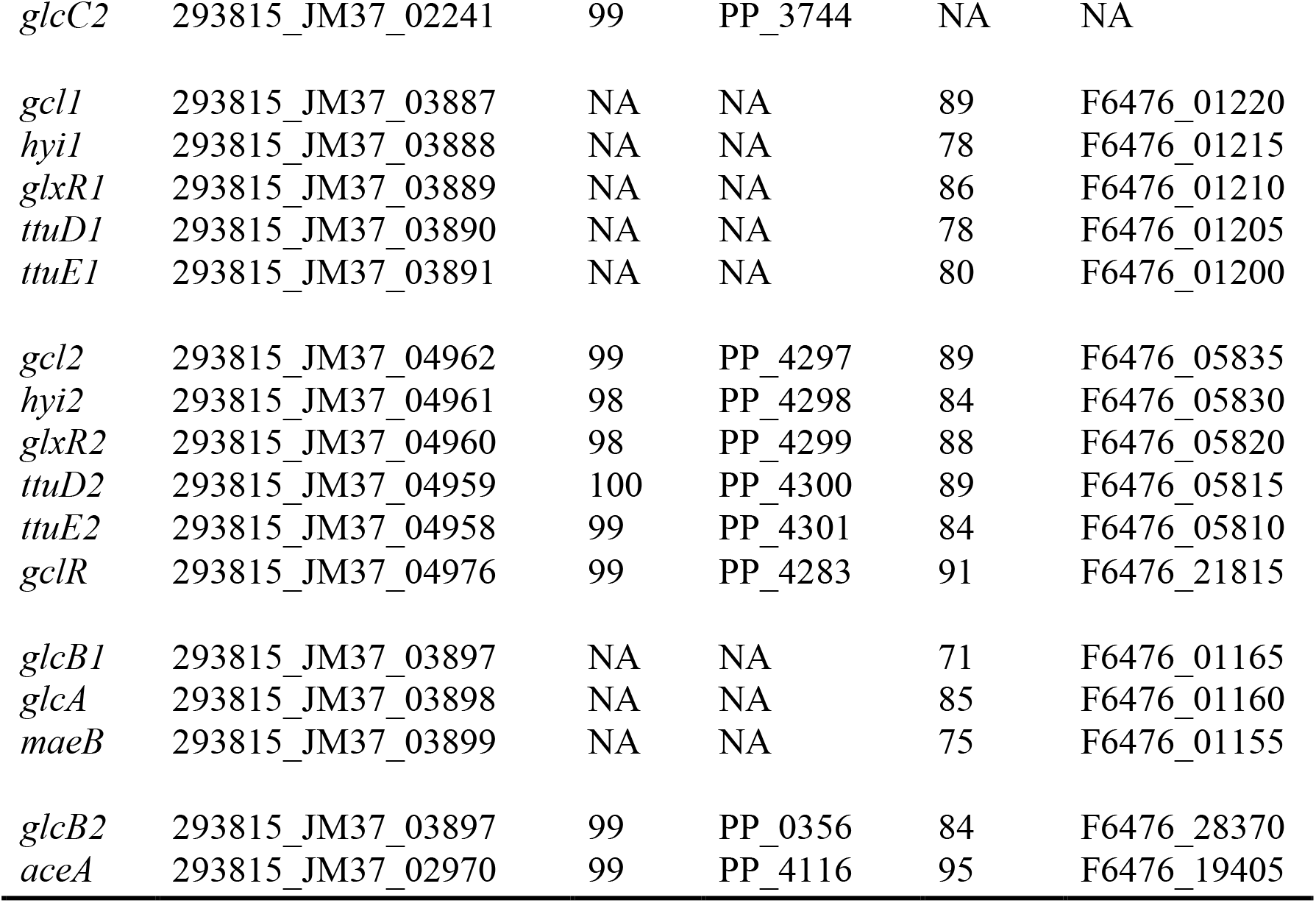
Comparation of the genes involved in EG metabolism present in *P. putida* JM37 and their homologous in *P. putida* KT2440 and *P. unsongensis* GO16. Results are expressed in percentage of amino acid identity.

The genome of *P. putida* JM37 contains both a *gclR* copy and the GclR putative binding sites upstream *gcl* present in KT2440 (Novichkov et al., 2013), making intriguing why this strain can grow on EG but *P. putida* KT2440 not. This is probably due to the presence in *P. putida* JM37 of an operon containing the genes necessary for transforming glycolate to pyruvate, i.e, *gcl, hyi, glxR, ttuD, ttuE, glcDEF* and *glcB*, and also a glycolate permease (*glcA*) and a malic enzyme (*maeB*), which could lead malate to pyruvate. In this case, the region upstream *gcl* does not contain the putative binding sites of GclR, so it should not be repressed by this regulator (Fig. S1).

In *P. putida* JM37, both copies of *glcDEF* were accompanied by *glcC* and *glcG*. In *E. coli*, inactivation of *glcC* made both GlcDEF and GlcB activity undetectable, whereas when this gen was transformed in a plasmid into a Δ*glcC* strain, both GlcDEF and GlcB restored their activities. This suggest that GlcC acts as an activator for these enzymes in *E. coli*. On the other hand, *glcG* function is not clear, as its disruption does not suppose any change in GlcDEF nor GlcB activities (Pellicer et al., 1996). However, in *P. putida* KT2440 *glcG* was upregulated alongside *glcDEF* when growing on glycerol in the presence of lanthanum (Wehrmann et al., 2020).

The two copies of *glcC* present in *P. putida* JM37 may improve GlcDEF activity. We suggest that the natural ability of *P. putida* JM37 to efficiently grow on EG is due to its genetic redundance. First, the two *glcCDEFG* copies would allow the bacterium to surpass the glycolaldehyde accumulation bottleneck and get more glyoxylate. Second, the non-repressed Gcl pathway and the second *glcB* alongside *maeB* would increase biomass formation from glyoxylate.

The *P. denitrificans* genes corresponding to the β-hydroxyaspartate cycle (*bhcCBD*) were not found in *P. putida* JM37, discarding this EG metabolization pathway.

### Disruption of the Gcl Pathway and the Glyoxylate Shunt Prevents P. putida JM37 to Grow on EG

In a proteomic study, Mückschel et al. (2012) observed an up-regulation of AceA, Gcl and GlcB in *P. putida* JM37 in presence of EG, suggesting that this strain may employ both the Gcl pathway and the glyoxylate shunt. However, disruption of *aceA, gcl* and *glcB* did not interrupt glyoxylate consumption in JM37, suggesting that this bacterium may use an alternative pathway for EG assimilation (Li et al., 2010). Given the *glcDEF* and Gcl pathway operon duplication found in JM37’s genome, we hypothesise that the in the previous work only one of the two copies of the *glcB* and *gcl* genes were disrupted. To validate this hypothesis we disrupted the five key genes for the Gcl pathway and the glyoxylate shunt present in JM37 (*gcl* x2, *glcB* x2 and *aceA* x2) employing a CRISPR-dCas9 base editing method (Volke et al., 2022).

The mutant strain *P. putida* JM37-ΔEG was not able to grow on 10 mM EG, while the wild-type strain reached the stationary phase at 0.6 OD^600^ after 24 h (Fig. S2). The mutant strain grew normally in LB, which suggests that potential off-target mutations did not result in a significant fitness defect.

Furthermore, the genome of *P. putida* JM37-ΔEG was sequenced, and the target mutations were confirmed. We found collateral C**→**T substitutions inside the target region in 4/5 targeted genes inside the gRNA region, but none interfered with the insertion of the intended early STOP codons (Fig. S3).

Our results support the hypothesis that *P. putida* JM37 employs the EG assimilation pathway present in *P. putida* KT2440 and *P. umsongensis* GO16. This experiment also validates the use of the CRISPR-Cas9 base editor (Volke et al. (2022)) in *P. putida* JM37, which could facilitate future research in this bacterium. The gRNAs employed and the native and mutant aligned sequences are shown in Table S1.

### P. putida JM37 Accumulates mcl-PHA Using EG as a Growth Substrate

Besides biomass generation, we evaluated *P. putida* JM37 capacity to upcycle EG to mcl-PHAs. Prior quantification, the presence of PHA granules was confirmed by Nile Red staining (Fig. 4). *P. putida* JM37 accumulated 15.68 ± 1.63 % PHA/cell dry weight (CDW) in 24 h growing in shake flasks with 50 mM EG. The nitrogen source (NH_4_Cl) was limited to 0.1 g/L (C/N = 53.5), as greater nitrogen concentrations resulted in lower or no presence of granules, confirmed by Nile Red staining. The mcl-PHA profile was composed by C6 (2.28 ± 0.07 %), C8 (12.36 ± 0.54 %), C10 (60.16 ± 0.97 %), C12:1 (8.32 ± 0.19 %) and C12 (10.85 ± 0.22 %) (Fig. 4).

**Figure 4.**
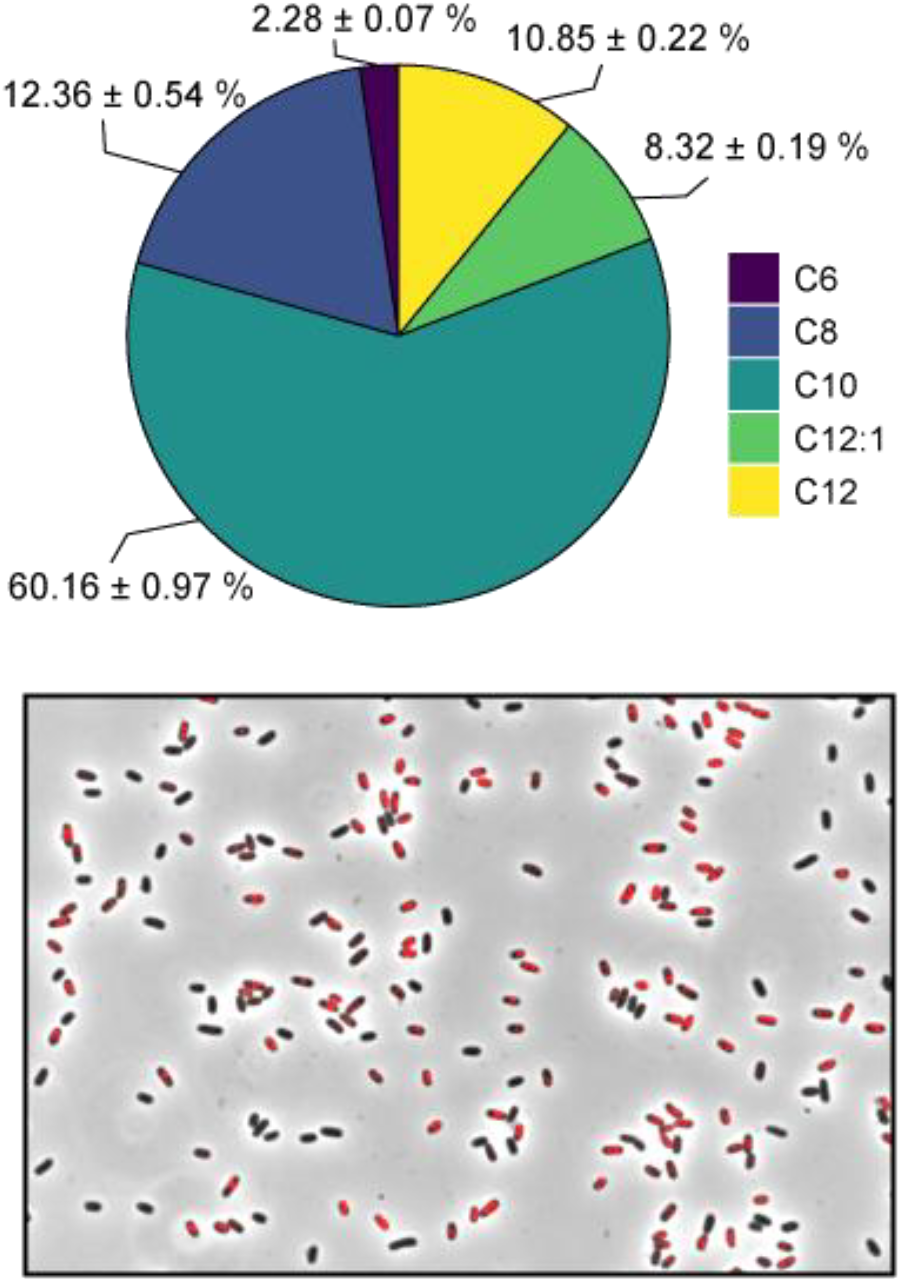
Upper panel: PHA profile of *P. putida* JM37 using 50 mM EG as the sole carbon source after 24 h (total PHA/CDW of 15.68 ± 1.63 %, with ∼60% of C10). Lowe panel: Nile Red staining of *P. putida* JM37 growing on 50 mM EG after 24 h (overlayed red and phase contrast channels).

In *P. putida*, nitrogen starvation is known to increase the conversion of *de novo* intermediates, i.e, those not related structurally to PHA, such as EG, into PHA monomers (Prieto et al., 2016). Under nitrogen limitation (C/N = 100), *P. putida* KT2440 was able to accumulate 32.19 % PHA/CDW employing 100 mM EG in 72 h, with similar accumulation rates using acetate as carbon source (Franden et al., 2018). *P. umsongensis* GO16 also accumulated mcl-PHA when growing on substrates not-related to PHA, although not with EG as the sole carbon source (Narancic et al., 2021). The evolved strain *P. umsongensis* GO16 KS3 was able to accumulate 7% PHD/CDW in a 5 L bioreactor with a mixture of EG and TPA, mainly composed of C10, C8 and C12 with a predominance of C10 (Tiso et al., 2021), similarly to *P. putida* JM37. This composition profile is common in *P. putida* when employing PHA structurally not-related substrates, such as glucose (Borrero-de Acuña et al., 2014), glycerol (Escapa et al., 2013), *p*-coumaric acid (Salvachúa et al., 2020) or ethanol (De Vrije et al., 2023). Furthermore, genetic modifications have been shown to increase PHA accumulation in *Pseudomonas* sp. The overexpression of the pyruvate dehydrogenase subunit gene *acoA* alongside the deletion of the glucose dehydrogenase gene *gcd* resulted in a 120% increase of PHA production from glucose in *P. putida* KT2440 (Borrero-de Acuña et al., 2014). Based on metabolic model predictions, Manoli et al. (2024) performed a series of knock-outs in *P. putida* KT2440 targeting various metabolic pathways, finding particularly important the deletion of succinyl-CoA synthase (*sucC, D*) and isocitrate lyase (*aceA*), which uncoupled nitrogen limitation to PHA production and increased its accumulation up to 33.7% of PHA/CDW (∼ 4.75-fold increase compared to wild-type) employing 4-hydroxybenzoate and octanoate as carbon sources.

In addition, it is interesting to note that *P. putida* JM37 can only partially upcycle PET degradation products, as it cannot assimilate TPA. This problem has been addressed in *P. putida* KT2440 by cloning the *tph* degradation cluster, making it able to assimilate TPA (Manoli et al., 2024; Hara et al., 2007; Werner et al., 2021). On the other hand, an interesting option for a complete PET upcycling is the design of artificial consortia, in which one strain is able to consume EG and the other TPA. This strategy has been recently tested in with *P. putida*, comparing the efficiency of a strain designed to assimilate both monomers with a consortium formed by two strains, each specialized in either EG or TPA. The consortium assimilating a mixture of monomers grew faster and achieved 92% more mcl-PHA and 2.82-fold higher *cis*-*cis* muconate production compared to the single strain (Bao et al., 2023).

### P. putida JM37 Expresses Heterologous Biosynthetic Pathways

Characteristics such as rapid grow, versatile metabolism, and capacity to express heterologous pathways make *P. putida* a common chassis in synthetic biology (Nikel & De Lorenzo, 2018). To use *P. putida* JM37 as a chassis for developing synthetic biology strategies in white biotechnology, as a proof of concept we tested this bacterium’s capacity to produce the pigment violacein.

Violacein was chosen due to its detection simplicity and relative complex synthesis, as its biosynthetic cluster is composed of five enzymes, VioABCDE, (Zhang et al., 2025). This pathway yields a combination of violacein and deoxyviolacein, which is known as “crude violacein” (Zhang et al., 2025). *P. putida* Vi crude violacein production was confirmed visually in an agar plate (Fig. S4). Furthermore, this strain was able to produce 12.9 ± 1.6 mg/L of crude violacein in 18 h in shake flasks employing 50 mM EG as the sole carbon source.

Several studies have used *vioABCDE* to test the capabilities of different bacteria on various substrates. For example, *P. putida* KT2440 was used to produce violacein using a nylon hydrolysate, obtaining 38 mg/L in 65 h (de Witt et al., 2025). *Pseudoalteromonas atlantica* T6c was used to produce violacein using red algal biomass as substrate, obtaining 34 µg/mL with agaropectin as carbon source (Pathiraja et al., 2023). High yields of violacein production can be achieved employing conventional carbon sources such as glucose, galactose or glycerol, as well as nutrient rich agricultural byproducts. Also, addition of L-tryptophan helps to increase yields, as this amino acid is a precursor for violacein biosynthesis (Zhang et al., 2025). To the best of our knowledge, this is the first work that reports crude violacein production from EG.

### P. putida JM37 Expresses and Secretes Heterologous PET-Hydrolases

As PET is a source of EG, we transformed the PET-degrading enzyme FAST-PETase to assess the ability of *P. putida* JM37 to produce and secrete heterologous enzymes.

PETase strains, *P. putida* 3-FP and *P. putida* 5-FP, were confirmed to produce and secrete active FAST-PETase to the extracellular medium by *p*NPB hydrolysis. Activity values reached 0.36 ± 0.05 U/mL for *P. putida* 3-FP and 0.41 ± 0.006 U/mL for *P. putida* 5-FP. Wild-type and non-induced strains were used as negative controls, confirming negligible activity against *p*NPB (Fig. 5). The enzyme was secreted employing a SPpstu signal peptide from *P. stutzeri* MO19, which belongs to the Sec/SPI secretion system (Teufel et al., 2022). In addition, Tat secretion system genes (*tatA, tatB, tatC*) were found in *P. putida* JM37’s genome.

**Figure 5.**
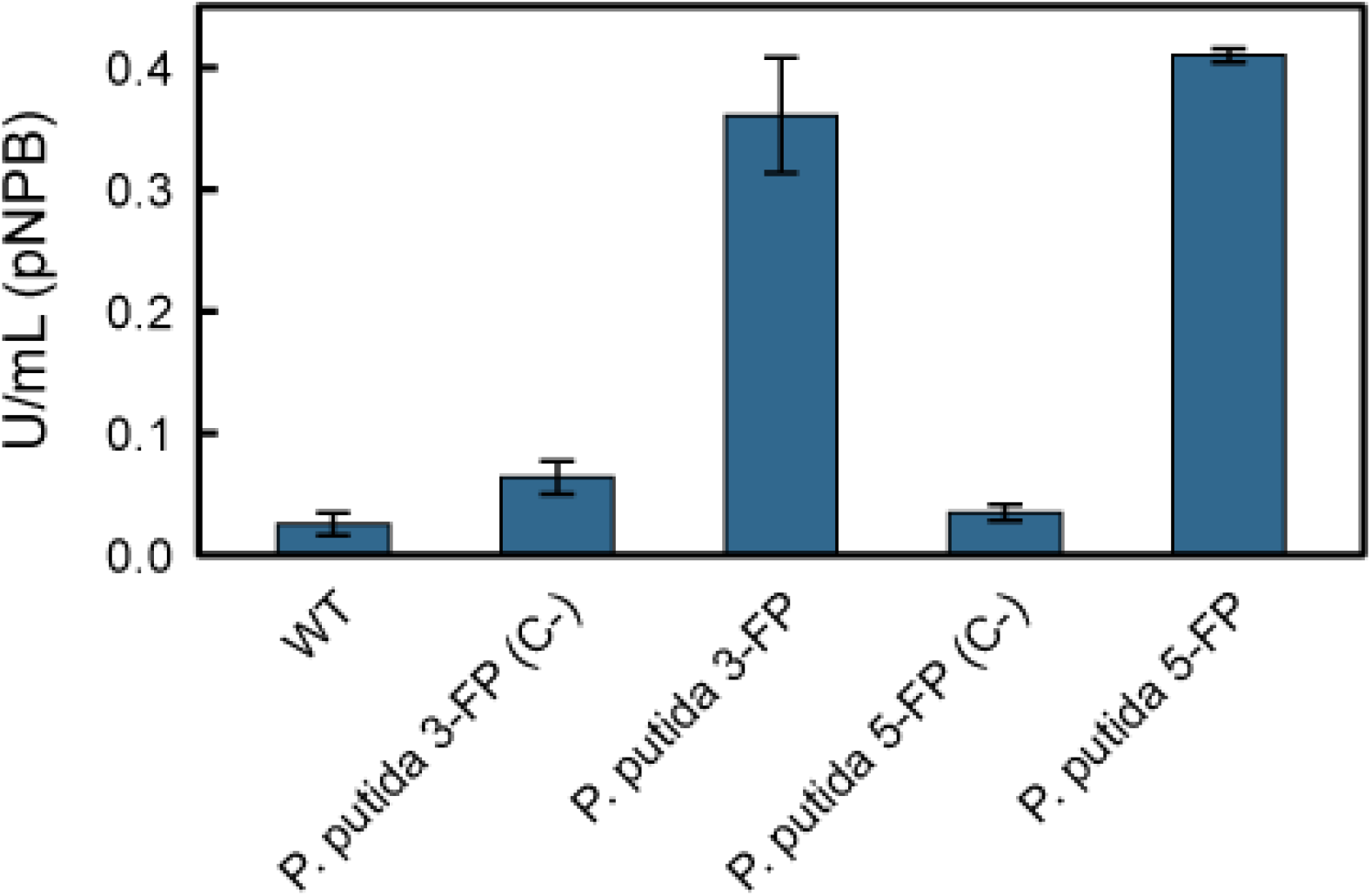
Recombinant *P*. putida JM37 strains’ activity against *p*NPB (U/mL). The results show a little advantage of *P. putida* 5-FP strain, as well as a negligible activity of the wild-type strain and little leaking in the non-induced controls.

Several works have employed *P. putida* for PET-hydrolase production. *P. putida* KT2440 was used as a host for FAST-PETase extracellular expression, as well as for *I. sakaiensis* PETase (*Is*PETase) and modified versions such as DuraPETase and ThermoPETase (Lu et al., 2022). Moreover, Brandenberg et al. (2022) expressed three well known PET hydrolases (LCC, HiC and *Is*PETase) in *P. putida* KT2440 trying different combinations involving genomic or plasmid expression, different promoters, and signal peptides. Here we demonstrated the functional expression of the FAST-PETase in *P. putida* JM37, making it a suitable host for the production and secretion of heterologous proteins.

## Conclusions

In this work, we propose *Pseudomonas putida* JM37 is a microbial platform for EG upcycling. Our genomic study, supported by CRISPR-cas9 driven knock outs, revealed a genetic redundance in glyoxylate metabolism that explains its robust growth on EG.

Beyond biomass formation, JM37 efficiently channels EG into medium-chain polyhydroxyalkanoates, revealing its potential for sustainable bioplastic production. We further established *P. putida* JM37 as a synthetic biology chassis by validating its capacity to express complex heterologous pathways. This includes the production of crude violacein directly from EG and the secretion of an active PET-hydrolase.

Together, these findings highlight *P. putida* JM37 as a bacterial chassis that couples efficient EG metabolism with biotechnological versatility. Its natural metabolic capabilities, combined with tractable genetic tools, make JM37 a promising candidate for future strategies in PET waste upcycling and the sustainable biosynthesis of value-added compounds.

## Acknowledgements

We sincerely thank Dr. Bernhard Hauer (University of Stuttgart, Germany) for kindly providing us with the *Pseudomonas putida* JM37 strain. This work was supported by the Spanish projects MICODE (PID2020-114210RB-I00 MCIN/AEI), DEMO (MCIN/AEI/”NextGenerationEU”/PRTR, TED2021-130096B-I00) and MOLA (PID2024-162673NB-I00 MCIN/AEI). The authors would also like to thank IBISBA1.0 project (H2020 730976), the SusPlast-CSIC Interdisciplinary Platform, and the LifeHub-CSIC research network for their support. J. Jiménez acknowledges the support received from the Biotechnology and Biological Sciences Research Council (BBSRC) through grants BB/Y007972/1 and BB/T011289/1 from the ERA-CoBioTech programme of the EU.

## References

Aziz, R. K., Bartels, D., Best, A. A., DeJongh, M., Disz, T., Edwards, R. A., Formsma, K., Gerdes, S., Glass, E. M., Kubal, M., Meyer, F., Olsen, G. J., Olson, R., Osterman, A. L., Overbeek, R. A., McNeil, L. K., Paarmann, D., Paczian, T., Parrello, B., … Zagnitko, O. (2008). The RAST Server: Rapid Annotations using Subsystems Technology. BMC Genomics, 9(1), 75. 10.1186/1471-2164-9-75

Barragán, M. J. L., Carmona, M., Zamarro, M. T., Thiele, B., Boll, M., Fuchs, G., García, J. L., & Díaz, E. (2004). The bzd Gene Cluster, Coding for Anaerobic Benzoate Catabolism, in Azoarcus sp. Strain CIB. Journal of Bacteriology, 186(17), 5762–5774. 10.1128/JB.186.17.5762-5774.2004

Blin, K., Pedersen, L. E., Weber, T., & Lee, S. Y. (2016). CRISPy-web: An online resource to design sgRNAs for CRISPR applications. Synthetic and Systems Biotechnology, 1(2), 118–121. 10.1016/j.synbio.2016.01.003

Boronat, A., Caballero, E., & Aguilar, J. (1983). Experimental evolution of a metabolic pathway for ethylene glycol utilization by Escherichia coli. Journal of Bacteriology, 153(1), 134–139. 10.1128/jb.153.1.134-139.1983

Borrero-de Acuña, J. M., Bielecka, A., Häussler, S., Schobert, M., Jahn, M., Wittmann, C., Jahn, D., & Poblete-Castro, I. (2014). Production of medium chain length polyhydroxyalkanoate in metabolic flux optimized Pseudomonas putida. Microbial Cell Factories, 13(1), 88. 10.1186/1475-2859-13-88

Brandenberg, O. F., Schubert, O. T., & Kruglyak, L. (2022). Towards synthetic PETtrophy: Engineering Pseudomonas putida for concurrent polyethylene terephthalate (PET) monomer metabolism and PET hydrolase expression. Microbial Cell Factories, 21(1), 119. 10.1186/s12934-022-01849-7

Caballero, E., Baldomá, L., Ros, J., Boronat, A., & Aguilar, J. (1983). Identification of lactaldehyde dehydrogenase and glycolaldehyde dehydrogenase as functions of the same protein in Escherichia coli. Journal of Biological Chemistry, 258(12), 7788–7792. 10.1016/S0021-9258(18)32248-8

Carniel, A., Ferreira Dos Santos, N., Buarque, F. S., Mendes Resende, J. V., Ribeiro, B. D., Marrucho, I. M., Coelho, M. A. Z., & Castro, A. M. (2024). From trash to cash: Current strategies for bio-upcycling of recaptured monomeric building blocks from poly(ethylene terephthalate) (PET) waste. Green Chemistry, 26(10), 5708–5743. 10.1039/D4GC00528G

Carniel, A., Santos, A. G., Chinelatto, L. S., Castro, A. M., & Coelho, M. A. Z. (2023). Biotransformation of ethylene glycol to glycolic acid by Yarrowia lipolytica: A route for poly(ethylene terephthalate) (PET) upcycling. Biotechnology Journal, 18(6), 2200521. 10.1002/biot.202200521

Chen, S., Zhou, Y., Chen, Y., & Gu, J. (2018). fastp: An ultra-fast all-in-one FASTQ preprocessor. Bioinformatics, 34(17), i884–i890. 10.1093/bioinformatics/bty560

De Vrije, T., Nagtegaal, R. M., Veloo, R. M., Kappen, F. H. J., & De Wolf, F. A. (2023). Medium chain length polyhydroxyalkanoate produced from ethanol by Pseudomonas putida grown in liquid obtained from acidogenic digestion of organic municipal solid waste. Bioresource Technology, 375, 128825. 10.1016/j.biortech.2023.128825

de Witt, J., Luthe, T., Wiechert, J., Jensen, K., Polen, T., Wirtz, A., Thies, S., Frunzke, J., Wynands, B., & Wierckx, N. (2025). Upcycling of polyamides through chemical hydrolysis and engineered Pseudomonas putida. Nature Microbiology, 10(3), 667–680. 10.1038/s41564-025-01929-5

Diao, J., Hu, Y., Tian, Y., Carr, R., & Moon, T. S. (2023). Upcycling of poly(ethylene terephthalate) to produce high-value bio-products. Cell Reports, 42(1), 111908. 10.1016/j.celrep.2022.111908

Durante-Rodríguez, G., De Lorenzo, V., & Martínez-García, E. (2014). The Standard European Vector Architecture (SEVA) Plasmid Toolkit. In A. Filloux & J.-L. Ramos (Eds), Pseudomonas Methods and Protocols (Vol. 1149, pp. 469–478). Springer New York. 10.1007/978-1-4939-0473-0_36

Escapa, I. F., Del Cerro, C., García, J. L., & Prieto, M. A. (2013). The role of GlpR repressor in Pseudomonas putida KT2440 growth and PHA production from glycerol. Environmental Microbiology, 15(1), 93–110. 10.1111/j.1462-2920.2012.02790.x

Franden, M. A., Jayakody, L. N., Li, W.-J., Wagner, N. J., Cleveland, N. S., Michener, W. E., Hauer, B., Blank, L. M., Wierckx, N., Klebensberger, J., & Beckham, G. T. (2018). Engineering Pseudomonas putida KT2440 for efficient ethylene glycol utilization. Metabolic Engineering, 48, 197–207. 10.1016/j.ymben.2018.06.003

Fujita, M., Torigoe, K., Nakada, T., Tsusaki, K., Kubota, M., Sakai, S., & Tsujisaka, Y. (1989). Cloning and nucleotide sequence of the gene (amyP) for maltotetraose-forming amylase from Pseudomonas stutzeri MO-19. Journal of Bacteriology, 171(3), 1333–1339. 10.1128/jb.171.3.1333-1339.1989

Hachisuka, S., Chong, J. F., Fujiwara, T., Takayama, A., Kawakami, Y., & Yoshida, S. (2022). Ethylene glycol metabolism in the poly(ethylene terephthalate)-degrading bacterium Ideonella sakaiensis. Applied Microbiology and Biotechnology, 106(23), 7867–7878. 10.1007/s00253-022-12244-y

Johnson, N. W., Valenzuela-Ortega, M., Thorpe, T. W., Era, Y., Kjeldsen, A., Mulholland, K., & Wallace, S. (2025). A biocompatible Lossen rearrangement in Escherichia coli. Nature Chemistry, 17(7), 1020–1026. 10.1038/s41557-025-01845-5

Kawai, F., Iizuka, R., & Kawabata, T. (2024). Engineered polyethylene terephthalate hydrolases: Perspectives and limits. Applied Microbiology and Biotechnology, 108(1), 404. 10.1007/s00253-024-13222-2

Kenny, S. T., Runic, J. N., Kaminsky, W., Woods, T., Babu, R. P., Keely, C. M., Blau, W., & O’Connor, K. E. (2008). Up-Cycling of PET (Polyethylene Terephthalate) to the Biodegradable Plastic PHA (Polyhydroxyalkanoate). Environmental Science & Technology, 42(20), 7696–7701. 10.1021/es801010e

Kim, H. T., Kim, J. K., Cha, H. G., Kang, M. J., Lee, H. S., Khang, T. U., Yun, E. J., Lee, D.-H., Song, B. K., Park, S. J., Joo, J. C., & Kim, K. H. (2019a). Biological Valorization of Poly(ethylene terephthalate) Monomers for Upcycling Waste PET. ACS Sustainable Chemistry & Engineering, 7(24), 19396–19406. 10.1021/acssuschemeng.9b03908

Kim, H. T., Kim, J. K., Cha, H. G., Kang, M. J., Lee, H. S., Khang, T. U., Yun, E. J., Lee, D.-H., Song, B. K., Park, S. J., Joo, J. C., & Kim, K. H. (2019b). Biological Valorization of Poly(ethylene terephthalate) Monomers for Upcycling Waste PET. ACS Sustainable Chemistry & Engineering, 7(24), 19396–19406. 10.1021/acssuschemeng.9b03908

Li, W., Jayakody, L. N., Franden, M. A., Wehrmann, M., Daun, T., Hauer, B., Blank, L. M., Beckham, G. T., Klebensberger, J., & Wierckx, N. (2019). Laboratory evolution reveals the metabolic and regulatory basis of ethylene glycol metabolism by Pseudomonas putida KT2440. Environmental Microbiology, 21(10), 3669–3682. 10.1111/1462-2920.14703

Li, X. Z., Klebensberger, J. & Rosche, B. (2010). Effect of gcl, glcB and aceA Disruption on Glyoxylate Conversion by Pseudomonas putida JM37. Journal of Microbiology and Biotechnology, 20(6), 1006–1010. 10.4014/jmb.0912.12005

Lu, H., Diaz, D. J., Czarnecki, N. J., Zhu, C., Kim, W., Shroff, R., Acosta, D. J., Alexander, B. R., Cole, H. O., Zhang, Y., Lynd, N. A., Ellington, A. D., & Alper, H. S. (2022). Machine learning-aided engineering of hydrolases for PET depolymerization. Nature, 604(7907), 662–667. 10.1038/s41586-022-04599-z

Madeira, F., Madhusoodanan, N., Lee, J., Eusebi, A., Niewielska, A., Tivey, A. R. N., Lopez, R., & Butcher, S. (2024). The EMBL-EBI Job Dispatcher sequence analysis tools framework in 2024. Nucleic Acids Research, 52(W1), W521–W525. 10.1093/nar/gkae241

Manoli, M.-T., Gargantilla-Becerra, Á., del Cerro Sánchez, C., Rivero-Buceta, V., Prieto, M. A., & Nogales, J. (2024). A model-driven approach to upcycling recalcitrant feedstocks in Pseudomonas putida by decoupling PHA production from nutrient limitation. Cell Reports, 43(4), 113979. 10.1016/j.celrep.2024.113979

Mückschel, B., Simon, O., Klebensberger, J., Graf, N., Rosche, B., Altenbuchner, J., Pfannstiel, J., Huber, A., & Hauer, B. (2012). Ethylene Glycol Metabolism by Pseudomonas putida. Applied and Environmental Microbiology, 78(24), 8531–8539. 10.1128/AEM.02062-12

Narancic, T., Salvador, M., Hughes, G. M., Beagan, N., Abdulmutalib, U., Kenny, S. T., Wu, H., Saccomanno, M., Um, J., O’Connor, K. E., & Jiménez, J. I. (2021). Genome analysis of the metabolically versatile Pseudomonas umsongensis GO16: The genetic basis for PET monomer upcycling into polyhydroxyalkanoates. Microbial Biotechnology, 14(6), 2463–2480. 10.1111/1751-7915.13712

Nikel, P. I., & De Lorenzo, V. (2018). Pseudomonas putida as a functional chassis for industrial biocatalysis: From native biochemistry to trans-metabolism. Metabolic Engineering, 50, 142–155. 10.1016/j.ymben.2018.05.005

Novichkov, P. S., Kazakov, A. E., Ravcheev, D. A., Leyn, S. A., Kovaleva, G. Y., Sutormin, R. A., Kazanov, M. D., Riehl, W., Arkin, A. P., Dubchak, I., & Rodionov, D. A. (2013). RegPrecise 3.0 – A resource for genome-scale exploration of transcriptional regulation in bacteria. BMC Genomics, 14(1), 745. 10.1186/1471-2164-14-745

OECD. (2022). Global Plastics Outlook: Policy Scenarios to 2060. OECD. 10.1787/aa1edf33-en

Pang, J., Zheng, M., Sun, R., Wang, A., Wang, X., & Zhang, T. (2016). Synthesis of ethylene glycol and terephthalic acid from biomass for producing PET. Green Chemistry, 18(2), 342–359. 10.1039/C5GC01771H

Pathiraja, D., Park, B., Kim, B., Stougaard, P., & Choi, I.-G. (2023). Constructing Marine Bacterial Metabolic Chassis for Potential Biorefinery of Red Algal Biomass and Agaropectin Wastes. ACS Synthetic Biology, 12(6), 1782–1793. 10.1021/acssynbio.3c00063

Pellicer, M. T., Badía, J., Aguilar, J., & Baldomà, L. (1996). glc locus of Escherichia coli: Characterization of genes encoding the subunits of glycolate oxidase and the glc regulator protein. Journal of Bacteriology, 178(7), 2051–2059. 10.1128/jb.178.7.2051-2059.1996

Prieto, A., Escapa, I. F., Martínez, V., Dinjaski, N., Herencias, C., de la Peña, F., Tarazona, N., & Revelles, O. (2016). A holistic view of polyhydroxyalkanoate metabolism in Pseudomonas putida. Environmental Microbiology, 18(2), 341–357. 10.1111/1462-2920.12760

Prjibelski, A., Antipov, D., Meleshko, D., Lapidus, A., & Korobeynikov, A. (2020). Using SPAdes De Novo Assembler. Current Protocols in Bioinformatics, 70(1), e102. 10.1002/cpbi.102

Ren, M., Li, D., Addison, H., Noteborn, W. E. M., Andeweg, E. H., Glatter, T., de Winde, J. H., Rebelein, J. G., Lamers, M. H., & Schada von Borzyskowski, L. (2025). NAD-dependent dehydrogenases enable efficient growth of Paracoccus denitrificans on the PET monomer ethylene glycol. Nature Communications, 16(1), 5845. 10.1038/s41467-025-61056-x

Sadler, J. C., & Wallace, S. (2021). Microbial synthesis of vanillin from waste poly(ethylene terephthalate). Green Chemistry, 23(13), 4665–4672. 10.1039/D1GC00931A

Salvachúa, D., Rydzak, T., Auwae, R., De Capite, A., Black, B. A., Bouvier, J. T., Cleveland, N. S., Elmore, J. R., Furches, A., Huenemann, J. D., Katahira, R., Michener, W. E., Peterson, D. J., Rohrer, H., Vardon, D. R., Beckham, G. T., & Guss, A. M. (2020). Metabolic engineering of Pseudomonas putida for increased polyhydroxyalkanoate production from lignin. Microbial Biotechnology, 13(1), 290–298. 10.1111/1751-7915.13481

Schada von Borzyskowski, L., Schulz-Mirbach, H., Troncoso Castellanos, M., Severi, F., Gómez-Coronado, P. A., Paczia, N., Glatter, T., Bar-Even, A., Lindner, S. N., & Erb, T. J. (2023). Implementation of the β-hydroxyaspartate cycle increases growth performance of Pseudomonas putida on the PET monomer ethylene glycol. Metabolic Engineering, 76, 97–109. 10.1016/j.ymben.2023.01.011

Schuster, L. A., & Reisch, C. R. (2021). A plasmid toolbox for controlled gene expression across the Proteobacteria. Nucleic Acids Research, 49(12), 7189–7202. 10.1093/nar/gkab496

Shehata, W. M., Nady, T. G., Gad, F. K., Shoaib, A. M., & Bhran, A. A. (2024). A comparative study of mono ethylene glycol economic production via different techniques. Scientific Reports, 14(1), 28375. 10.1038/s41598-024-77713-y

Shimizu, T., Suzuki, K., & Inui, M. (2024). A mycofactocin-associated dehydrogenase is essential for ethylene glycol metabolism by Rhodococcus jostii RHA1. Applied Microbiology and Biotechnology, 108(1), 58. 10.1007/s00253-023-12966-7

Teufel, F., Almagro Armenteros, J. J., Johansen, A. R., Gíslason, M. H., Pihl, S. I., Tsirigos, K. D., Winther, O., Brunak, S., Von Heijne, G., & Nielsen, H. (2022). SignalP 6.0 predicts all five types of signal peptides using protein language models. Nature Biotechnology, 40(7), 1023–1025. 10.1038/s41587-021-01156-3

Tiso, T., Narancic, T., Wei, R., Pollet, E., Beagan, N., Schröder, K., Honak, A., Jiang, M., Kenny, S. T., Wierckx, N., Perrin, R., Avérous, L., Zimmermann, W., O’Connor, K., & Blank, L. M. (2021). Towards bio-upcycling of polyethylene terephthalate. Metabolic Engineering, 66, 167–178. 10.1016/j.ymben.2021.03.011

Trifunovic, D., Schuchmann, K., & Müller, V. (2016). Ethylene Glycol Metabolism in the Acetogen Acetobacterium woodii. Journal of Bacteriology, 198(7), 1058–1065. 10.1128/JB.00942-15

Volke, D. C., Martino, R. A., Kozaeva, E., Smania, A. M., & Nikel, P. I. (2022). Modular (de)construction of complex bacterial phenotypes by CRISPR/nCas9-assisted, multiplex cytidine base-editing. Nature Communications, 13(1), 3026. 10.1038/s41467-022-30780-z

Wang, H., Jiang, P., Lu, Y., Ruan, Z., Jiang, R., Xing, X.-H., Lou, K., & Wei, D. (2009). Optimization of culture conditions for violacein production by a new strain of Duganella sp. B2. Biochemical Engineering Journal, 44(2–3), 119–124. 10.1016/j.bej.2008.11.008

Wehrmann, M., Toussaint, M., Pfannstiel, J., Billard, P., & Klebensberger, J. (2020). The Cellular Response to Lanthanum Is Substrate Specific and Reveals a Novel Route for Glycerol Metabolism in Pseudomonas putida KT2440. mBio, 11(2), 10.1128/mbio.00516-20. 10.1128/mbio.00516-20

Wiegant, W. M., & De Bont, J. A. M. (1980). A New Route for Ethylene Glycol Metabolism in Mycobacterium E44. Microbiology, 120(2), 325–331. 10.1099/00221287-120-2-325

Zhang, Q., Yang, J., Mou, L., Jiang, Y., Barriuso, J., Guo, F., Xin, F., & Jiang, M. (2025). A comprehensive review on violacein production by microbial fermentation. BioDesign Research, 7(3), 100043. 10.1016/j.bidere.2025.100043

Zuriani, R., Vigneswari, S., Azizan, M. N. M., Majid, M. I. A., & Amirul, A. A. (2013). A high throughput Nile red fluorescence method for rapid quantification of intracellular bacterial polyhydroxyalkanoates. Biotechnology and Bioprocess Engineering, 18(3), 472–478. 10.1007/s12257-012-0607-z

