## Supplementary Material for "*Pseudomonas putida* JM37 as a novel bacterial chassis for ethylene glycol upcycling"

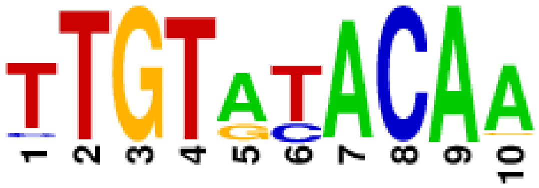


>gcl1_ KT2440

aaatagaacacaacgagcgatattttgtatacaatattttgaaacgatcgtatgcgatggcgcaaacctccgttttcacgggcgc
tctgacgaaagccagcttagccgatgaaaacccattgacctaagccgtcaggcgtgaatacactctgtcgcaaagcaagttgt
atacaattacaaaatcgatgaggcacaaaccATGAGCAAAATGAGAGCAATCGATG […]

>gcl2_JM

aaatagaacacaacgagatgtattttgtatacaatattttgaaacgatcgtattcgatggcgcaaatagccgttttcacggccgct
ttgacgaaagcgagcccacccgatgaaaacccattgacctgagccgccaggcgtgaatacactctgtcgcaaagcaagttgt
atacaattacaaaatcgatgaggcacaaaccATGAGCAAAATGAGAGCAATCGATG […]

>gcl2_GO16

aaatagaacacaatgaattatttttttgtatacaataattctcgaaaagcgccgcatgcgacgaaaagccacagtagaacgggct
tccgacgaatgaaacgagcgcttgagaaaatggattgacctgcgccgtccgccgtgaatacactctgcgcaaagccacttgta
tacaattacaaaacgtaaagaggcacaaaaccATGAGCAAAATGAGAGCAATCGAAG […]

>gcl1_JM

gggtcacccccttcgctcagcgcggcagaagcgaagctgcccttctgggcaacctgaacgaagctatgtagctctgtgtaccgt
gccattcgatacctttcatatcgacagaaaacatttttcaggtgttctaacgcacgcaagcctaatacagactgacctccactaca
aaaattatcgacccgtgcgaaggagtacacaATGGCCAAAATGAGAGCAATCGAGG […]

>gcl1_GO16

ggagtcaccccctcattcagtgcagcggcggcaaagctgcccttatgggcgacctgaataaaactgtgtagctcgacataacg
acccatcgatacctcatgtatcggcagaaaccgatttttggctattcacgccaataataacctgctccatgctgagatccgctaat
aaaaatatcgaccagcctgaaggaggcaataATGGCCAGAATGAGAGCAATCGAGG […]

**Figure S1.** GclR binding site predicted by RegPrecise (Novichkov et al., 2013). Sequences correspond to the beginning of *gcl* genes in *P. putida* KT2440, *P. putida* JM37 and *P. umsongensis* GO16. GclR binding sites are marked in yellow. Capital letters indicate *gcl*.

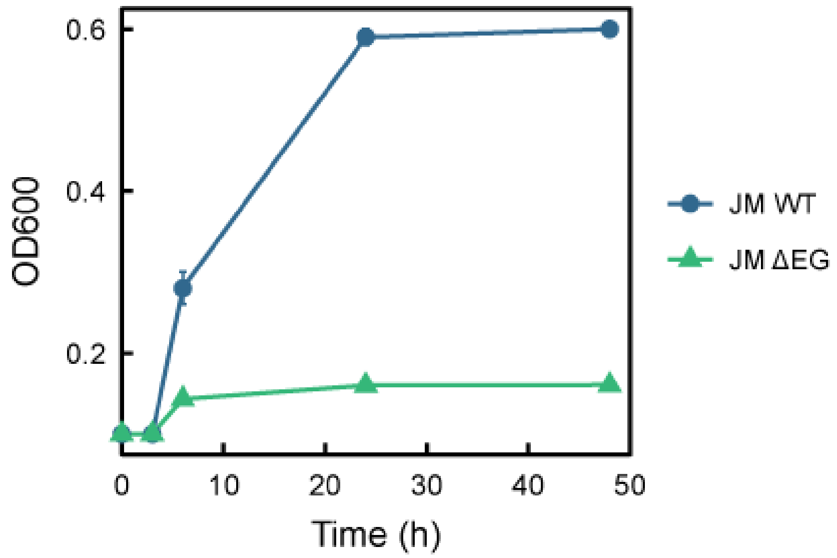


**Figure S2.** Growth of *P. putida* JM37 and *P. putida* JM37-ΔEG strains with EG 10 mM as the sole carbon source in shaking flasks cultures. Results are given as an average of n = 3.

**Table S1.** Primers employed for CRISPR gRNA design. Forward primers contain the gRNA (marked in yellow) obtained in CRISPy web service. Reverse primers are canonical, provided by Volke et al. (2022) supplementary material.

| **Target gene** | **Primer ID** | **Primer sequence** |
|---|---|---|
| *gcl1* | gcl1_Fw | atcgaggtctcCgtggCCCTCGAGCTCGACTGCACAGTTTTAGAGCTAGAAATAGC |
| | gRNA1-GG-R | atcgaggtctcCacctTTAGCTGCCTATACGGCAGT |
| *gcl2* | gcl2_Fw | atcgaggtctcCaggtGTGGTCGCCCACTTGGTCACGTTTTAGAGCTAGAAATAGC |
| | gRNA2-GG-R | atcgaggtctcCccgcTTAGCTGCCTATACGGCAGT |
| *glcB1* | glcB1_Fw | atcgaggtctcCttgtgaggcctatcagcgcttcctGTTTTAGAGCTAGAAATAGC |
| | gRNA3-GG-R | atcgaggtctcCaacaTTAGCTGCCTATACGGCAGT |
| *glcB2* | glcB2_Fw | atcgaggtctcCgcggcgttcaagtcggtggccttcGTTTTAGAGCTAGAAATAGC |
| | gRNA4-GG-R | atcgaggtctcCacaaTTAGCTGCCTATACGGCAGT |
| *aceA* | aceA_Fw | atcgaggtctcCtgttCCCTGACCAGTCGCTGTACCGTTTTAGAGCTAGAAATAGC |
| | gRNALastPosition-GG-R | ATCGAGGTCTCCAAACTTTCTTAGCTGCCTATACGG |

```
        1                                                50
gcl1_WT atggccaaaa tgagagcaat cgaggccgcc gttctggtgc tgcgccgtga
gcl1_EG ATGGCCAAAA TGAGAGCAAT CGAGGCCGCC GTTCTGGTGC TGCGCCGTGA

        51                                              100
gcl1_WT aggtatcgat accgcattcg gtatccccgg agccgcaatc aatccgttct
gcl1_EG AGGTATCGAT ACCGCATTCG GTATCCCCGG AGCCGCAATC AATCCGTTCT

        101                                             150
gcl1_WT acgctgcctt gaaaaaaatt ggtggaatcg accatgtatt ggcccgccat
gcl1_EG ACGCTGCCTT GAAAAAAATT GGTGGAATCG ACCATGTATT GGCCCGCCAT

        151                                             200
gcl1_WT gtcgagggtg cctcgcacat ggccgagggc tacacccgta cccacgcagg
gcl1_EG GTCGAGGGTG CCTCGCACAT GGCCGAGGGC TACACCCGTA CCCACGCAGG

        201                                             250
gcl1_WT taatatcggc ctgtgcatcg gcacatccgg cccggctggc accgacatgg
gcl1_EG TAATATCGGC CTGTGCATCG GCACATCCGG CCCGGCTGGC ACCGACATGG

        251                                             300
gcl1_WT tcactggcct gtacagcgcc tcggcagatt ccattccgat tctctgcatc
gcl1_EG TCACTGGCCT GTACAGCGCC TCGGCAGATT CCATTCCGAT TCTCTGCATC

        301                                             350
gcl1_WT acagggcaag cccctcgagc tcgactgcac aaggaagact tccaggccgt
gcl1_EG ACAGGGCAAG CCCTTTGAGC TTGACTGCAC AAGGAAGACT TCCAGGCCGT

        351                                             400
gcl1_WT cgacatcacc agcattgtca aaccggtgac caagtgggcc actactgtgc
gcl1_EG CGACATCACC AGCATTGTCA AACCGGTGAC CAAGTGGGCC ACTACTGTGC

        401                                             450
gcl1_WT tggagccggg tcaggtgccc tatgccttcc agaaagcttt ttacgaaatg
gcl1_EG TGGAGCCGGG TCAGGTGCCC TATGCCTTCC AGAAAGCTTT TTACGAAATG

        451                                             500
gcl1_WT cgcagtggcc ggcccggccc agtactgatc gacctgccat ttgacgtgca
gcl1_EG CGCAGTGGCC GGCCCGGCCC AGTACTGATC GACCTGCCAT TTGACGTGCA

        501                                             550
gcl1_WT gatggccgag atcgagtttg acatcgacgc ctacgagccg ctaccggtgc
gcl1_EG GATGGCCGAG ATCGAGTTTG ACATCGACGC CTACGAGCCG CTACCGGTGC

        551                                             600
gcl1_WT acaagcccca agccagtcgc gttcaggccg agaaggctct tgccatgctc
gcl1_EG ACAAGCCCCA AGCCAGTCGC GTTCAGGCCG AGAAGGCTCT TGCCATGCTC

        601                                             650
gcl1_WT aatgccgctg agcgcccgct gatcgtcgcc ggtggcggta tcatcaacgc
gcl1_EG AATGCCGCTG AGCGCCCGCT GATCGTCGCC GGTGGCGGTA TCATCAACGC

        651                                             700
gcl1_WT cgacgccagt gacaagctgg tggccttcgc cgaactgacc ggcgtgccgg
gcl1_EG CGACGCCAGT GACAAGCTGG TGGCCTTCGC CGAACTGACC GGCGTGCCGG

        701                                             750
gcl1_WT taatccctac cctgatgggc tggggaaccc tgcccgatga ccacccgttg
gcl1_EG TAATCCCTAC CCTGATGGGC TGGGGAACCC TGCCCGATGA CCACCCGTTG
```

```
        751                                                800
gcl1_WT atggcgggca tgtgtggcct gcagacctcg caccgctacg gcaacgctac
gcl1_EG ATGGCGGGCA TGTGTGGCCT GCAGACCTCG CACCGCTACG GCAACGCTAC

        801                                                850
gcl1_WT cctgctcgag tcggatctgg tgttcggcat cggcaatcgc tgggccaacc
gcl1_EG CCTGCTCGAG TCGGATCTGG TGTTCGGCAT CGGCAATCGC TGGGCCAACC

        851                                                900
gcl1_WT gccatactgg ctctgtcgac gtctacaccg aaggccgcac gtttgtgcac
gcl1_EG GCCATACTGG CTCTGTCGAC GTCTACACCG AAGGCCGCAC GTTTGTGCAC

        901                                                950
gcl1_WT gtggacatcg aaccgaccca gatcggtcgt gtgttcaccc cggacctcgg
gcl1_EG GTGGACATCG AACCGACCCA GATCGGTCGT GTGTTCACCC CGGACCTCGG

        951                                               1000
gcl1_WT catcgtctcc gacgcaggcg ccgccctcga tgtattcctc gaggtagctc
gcl1_EG CATCGTCTCC GACGCAGGCG CCGCCCTCGA TGTATTCCTC GAGGTAGCTC

        1001                                              1050
gcl1_WT gcgagtggca ggccgctggc aagctcaagg accgcaaagc ctgggctgaa
gcl1_EG GCGAGTGGCA GGCCGCTGGC AAGCTCAAGG ACCGCAAAGC CTGGGCTGAA

        1051                                              1100
gcl1_WT gcctgccgtg aacgcaaacg caccctgcaa cgcaagaccc acttcgacga
gcl1_EG GCCTGCCGTG AACGCAAACG CACCCTGCAA CGCAAGACCC ACTTCGACGA

        1101                                              1150
gcl1_WT agtgccggtc aagccgcaac gggtatacga ggaaatgaat cgtttcttcg
gcl1_EG AGTGCCGGTC AAGCCGCAAC GGGTATACGA GGAAATGAAT CGTTTCTTCG

        1151                                              1200
gcl1_WT gcaaggatac ttgctacgtc agtaccatcg gtctgtcgca gattgccggc
gcl1_EG GCAAGGATAC TTGCTACGTC AGTACCATCG GTCTGTCGCA GATTGCCGGC

        1201                                              1250
gcl1_WT gcccagttcc tgcacgtgta caagccgcgt cactggatca actgcggcca
gcl1_EG GCCCAGTTCC TGCACGTGTA CAAGCCGCGT CACTGGATCA ACTGCGGCCA

        1251                                              1300
gcl1_WT ggcggggccg ttgggttgga ccatccccgc tgctctcggc gtggtgaaag
gcl1_EG GGCGGGGCCG TTGGGTTGGA CCATCCCCGC TGCTCTCGGC GTGGTGAAAG

        1301                                              1350
gcl1_WT ccgatccaga gcgccaggtc gtcgcgttgt ctggtgacta cgacttccag
gcl1_EG CCGATCCAGA GCGCCAGGTC GTCGCGTTGT CTGGTGACTA CGACTTCCAG

        1351                                              1400
gcl1_WT ttcatgatcg aggaactggc ggtgggcgca cagttcaacc tgccgtacat
gcl1_EG TTCATGATCG AGGAACTGGC GGTGGGCGCA CAGTTCAACC TGCCGTACAT

        1401                                              1450
gcl1_WT ccacgtgctg gtgaataacg cctacctcgg tctgattcgc caagcccagc
gcl1_EG CCACGTGCTG GTGAATAACG CCTACCTCGG TCTGATTCGC CAAGCCCAGC

        1451                                              1500
gcl1_WT gtggttttga catcgactac tgcgtgcagc tgtcgttcga aaacgtcaac
```

```
gcl1_EG GTGGTTTTGA CATCGACTAC TGCGTGCAGC TGTCGTTCGA AAACGTCAAC

        1501                                            1550
gcl1_WT gcgccagaac tgggtagcta cggcgtggac cacgtgaccg tggttgaggg
gcl1_EG GCGCCAGAAC TGGGTAGCTA CGGCGTGGAC CACGTGACCG TGGTTGAGGG

        1551                                            1600
gcl1_WT cttaggctgc aaggccatcc gtgtattcga cgcgaaccag ttgcccgccg
gcl1_EG CTTAGGCTGC AAGGCCATCC GTGTATTCGA CGCGAACCAG TTGCCCGCCG

        1601                                            1650
gcl1_WT ccttcgccca ggcccgcgaa ctgatggaaa ccttccgcgt accggtggtg
gcl1_EG CCTTCGCCCA GGCCCGCGAA CTGATGGAAA CCTTCCGCGT ACCGGTGGTG

        1651                                            1700
gcl1_WT gtcgaggtaa tcctggagcg cgtcacgaac atcgccatgg gtactgagat
gcl1_EG GTCGAGGTAA TCCTGGAGCG CGTCACGAAC ATCGCCATGG GTACTGAGAT

        1701                                            1750
gcl1_WT caacgccatt aacgaatttg aggcgctggc taccagtcga gccgacgccc
gcl1_EG CAACGCCATT AACGAATTTG AGGCGCTGGC TACCAGTCGA GCCGACGCCC

        1751                    1776
gcl1_WT ctacttccat cctgccactg gactga
gcl1_EG CTACTTCCAT CCTGCCACTG GACTGA


        1                                                 50
gcl2_WT tcagtccagc agcgagatgg cggttggcgc gtcgttgccg accagggcca
gcl2_EG TCAGTCCAGC AGCGAGATGG CGGTTGGCGC GTCGTTGCCG ACCAGGGCCA

        51                                               100
gcl2_WT ggtcttcgaa ttcgttgacc gcgttgatct cggtgcccat ggaaatgttg
gcl2_EG GGTCTTCGAA TTCGTTGACC GCGTTGATCT CGGTGCCCAT GGAAATGTTG

        101                                              150
gcl2_WT gtcacacgct cgagaatcac ttcaaccacc accggcacgc ggaactcttc
gcl2_EG GTCACACGCT CGAGAATCAC TTCAACCACC ACCGGCACGC GGAACTCTTC

        151                                              200
gcl2_WT ggccatcttc tgtgccttga tcagggcagg ggcgatttct gctggctcga
gcl2_EG GGCCATCTTC TGTGCCTTGA TCAGGGCAGG GGCGATTTCT GCTGGCTCGA

        201                                              250
gcl2_WT acacacggat ggccttgcaa cccaggcctt cgaccacagc gacgtggtcg
gcl2_EG ACACACGGAT GGCCTTGCAA CCCAGGCCTT CGACCACAGC GACGTGGTCG

        251                                              300
gcl2_WT acaccgtagg tggcagcgtc ggtcgagttg atgttctcga acgccagttg
gcl2_EG ACACCGTAGG TGGCAGCGTC GGTCGAGTTG ATGTTCTCGA ACGCCAGTTG

        301                                              350
gcl2_WT tacacagtaa tccatgtcga agccacgctg cgcctggcgg atcaggccca
gcl2_EG TACACAGTAA TCCATGTCGA AGCCACGCTG CGCCTGGCGG ATCAGGCCCA

        351                                              400
gcl2_WT ggtaggcgtt gttcaccagt acgtgaacgt acggcaggtt gaactgcgca
gcl2_EG GGTAGGCGTT GTTCACCAGT ACGTGAACGT ACGGCAGGTT GAACTGCGCA
```

```
        401                                               450
gcl2_WT cccaccgcca gttcttcgat catgaactgg aagtcgtagt cacccgacag
gcl2_EG CCCACCGCCA GTTCTTCGAT CATGAACTGG AAGTCGTAGT CACCCGACAG

        451                                               500
gcl2_WT cgccacaacc ttgcgcttcg gatcggcttt gaccacgccc agtgcagcag
gcl2_EG CGCCACAACC TTGCGCTTCG GATCGGCTTT GACCACGCCC AGTGCAGCAG

        501                                               550
gcl2_WT ggatggtcca gcccagcggg ccggcctggc cgcagttgat ccagtggcgt
gcl2_EG GGATGGTCCA GCCCAGCGGG CCGGCCTGGC CGCAGTTGAT CCAGTGGCGT

        551                                               600
gcl2_WT ggcttgtaca catgcaggaa ctgcgcgccg gcgatctgcg acaggccgat
gcl2_EG GGCTTGTACA CATGCAGGAA CTGCGCGCCG GCGATCTGCG ACAGGCCGAT

        601                                               650
gcl2_WT ggtgctgacg tagcaggtgt ctttgccgaa cacctggttc atttcttcgt
gcl2_EG GGTGCTGACG TAGCAGGTGT CTTTGCCGAA CACCTGGTTC ATTTCTTCGT

        651                                               700
gcl2_WT agacgcgctg cggcttgacc ggcacgttgt cgaagtgggt cttgcgctgc
gcl2_EG AGACGCGCTG CGGCTTGACC GGCACGTTGT CGAAGTGGGT CTTGCGCTGC

        701                                               750
gcl2_WT aggctcgact tgcgctcctg gcactcttcc agccaggcct tgcggcactt
gcl2_EG AGGCTCGACT TGCGCTCCTG GCACTCTTCC AGCCAGGCCT TGCGGCACTT

        751                                               800
gcl2_WT gagcttgccg gcggctttcc actcgcgggc cacttccagg aacacgtcca
gcl2_EG GAGCTTGCCG GCGGCTTTCC ACTCGCGGGC CACTTCCAGG AACACGTCCA

        801                                               850
gcl2_WT gtgccttgcc agcatcggaa acgatgccca ggtccggggt gaacacgcgg
gcl2_EG GTGCCTTGCC AGCATCGGAA ACGATGCCCA GGTCCGGGGT GAACACGCGG

        851                                               900
gcl2_WT ccgatctggg tcggttcgat gtcgacgtgc acgaacttgc ggccttcggt
gcl2_EG CCGATCTGGG TCGGTTCGAT GTCGACGTGC ACGAACTTGC GGCCTTCGGT

        901                                               950
gcl2_WT gtagacatcg acggaaccgg tgtggcggtt ggcccagcgg ttaccgatac
gcl2_EG GTAGACATCG ACGGAACCGG TGTGGCGGTT GGCCCAGCGG TTACCGATAC

        951                                              1000
gcl2_WT cgaataccag gtcggacttc agcagggttg cgttgccata gcggtgcgag
gcl2_EG CGAATACCAG GTCGGACTTC AGCAGGGTTG CGTTGCCATA GCGGTGCGAG

        1001                                             1050
gcl2_WT gtctgcaggc caaccatgcc gaccatctgg gcgtggtcgt ccgggatggt
gcl2_EG GTCTGCAGGC CAACCATGCC GACCATCTGG GCGTGGTCGT CCGGGATGGT

        1051                                             1100
gcl2_WT gccccagccc atcagggtcg ggatcacggg tacgccggtc agttcggcga
gcl2_EG GCCCCAGCCC ATCAGGGTCG GGATCACGGG TACGCCGGTC AGTTCGGCGA

        1101                                             1150
gcl2_WT attcgaccag cttgtcgctg gcgtcggcgt tgatgatgcc gccaccggct
```

```
gcl2_EG ATTCGACCAG CTTGTCGCTG GCGTCGGCGT TGATGATGCC GCCACCGGCT

        1151                                             1200
gcl2_WT accagcagtg ggcgctcggc gtcattgagc agggccaggg ctttctcggc
gcl2_EG ACCAGCAGTG GGCGCTCGGC GTCATTGAGC AGGGCCAGGG CTTTCTCGGC

        1201                                             1250
gcl2_WT ctgtacgcgg gtggcggatg gcttgttgac tgccagtggt tcgtaggcat
gcl2_EG CTGTACGCGG GTGGCGGATG GCTTGTTGAC TGCCAGTGGT TCGTAGGCAT

        1251                                             1300
gcl2_WT cgatgtcgaa ttcgatttcg gccatctgca cgtcgaacgg caggtcgatc
gcl2_EG CGATGTCGAA TTCGATTTCG GCCATCTGCA CGTCGAACGG CAGGTCGATC

        1301                                             1350
gcl2_WT agcaccgggc ctgggcggcc ggtacgcatt tcataaaagg ccttctggaa
gcl2_EG AGCACCGGGC CTGGGCGGCC GGTACGCATT TCATAAAAGG CCTTCTGGAA

        1351                                             1400
gcl2_WT ggcgtaaggc acctggcctg gctccagaac ggtggtcgcc cacttggtca
gcl2_EG GGCGTAAGGC ACCTGGCCTG GCTCCAAAAC GGTGGTTGTT CACTTGGTCA

        1401                                             1450
gcl2_WT ctggcttgac gatgttggtg atgtcgacag cctggaagtc ttccttgtgc
gcl2_EG CTGGCTTGAC GATGTTGGTG ATGTCGACAG CCTGGAAGTC TTCCTTGTGC

        1451                                             1500
gcl2_WT aggcgggcac gtggcgcctg gccagtgatg cacagaatcg ggatggagtc
gcl2_EG AGGCGGGCAC GTGGCGCCTG GCCAGTGATG CACAGAATCG GGATGGAGTC

        1501                                             1550
gcl2_WT ggccgaggcg ctgtacaggc cggtgaccat gtcggtgccg gcagggccgg
gcl2_EG GGCCGAGGCG CTGTACAGGC CGGTGACCAT GTCGGTGCCG GCAGGGCCGG

        1551                                             1600
gcl2_WT aagtgccgat gcacacaccg atgttgcccg ggttggcgcg ggtgtagccc
gcl2_EG AAGTGCCGAT GCACACACCG ATGTTGCCCG GGTTGGCGCG GGTGTAGCCC

        1601                                             1650
gcl2_WT tcggccatgt gcgaggcacc ttcgacgtga cgagcgagga cgtgatcgat
gcl2_EG TCGGCCATGT GCGAGGCACC TTCGACGTGA CGAGCGAGGA CGTGATCGAT

        1651                                             1700
gcl2_WT gccaccgact ttcttcaggg ccgaatacaa cgggttgatg gcagcccccg
gcl2_EG GCCACCGACT TTCTTCAGGG CCGAATACAA CGGGTTGATG GCAGCCCCCG

        1701                                             1750
gcl2_WT ggatgccgaa cgcggtatct acaccttcac ggcgcatgac cagaacggct
gcl2_EG GGATGCCGAA CGCGGTATCT ACACCTTCAC GGCGCATGAC CAGAACGGCT

        1751                    1776
gcl2_WT gcatcgattg ctctcatttt gctcat
gcl2_EG GCATCGATTG CTCTCATTTT GCTCAT


         1                                                50
glcb1_WT atgaccgagc gcgtgccatg ccagcgtttg caggtggctg agaacttcaa
glcb1_EG ATGACCGAGC GCGTGCCATG CCAGCGTTTG CAGGTGGCTG AGAACTTCAA
```

```
           51                                                 100
glcb1_WT gcacttcatc gaggatgagg tgctccccgg caccgagatt acgtcggatc
glcb1_EG GCACTTCATC GAGGATGAGG TGCTCCCCGG CACCGAGATT ACGTCGGATC

           101                                                150
glcb1_WT atttctggag ggggctcgat gccctggtgc acgatctggc gccgcgcaac
glcb1_EG ATTTCTGGAG GGGGCTCGAT GCCCTGGTGC ACGATCTGGC GCCGCGCAAC

           151                                                200
glcb1_WT cgcgctttgc tggccgagcg tgttcacttg cagggcgaat tggatacctg
glcb1_EG CGCGCTTTGC TGGCCGAGCG TGTTCACTTG CAGGGCGAAT TGGATACCTG

           201                                                250
glcb1_WT gcaccgcgcc caccccgggc cggtaacgga catggaggcc tatcagcgct
glcb1_EG GCACCGCGCC CACCCCGGGC CGGTAACGGA CATGGAGGTT TATTAGCGCT

           251                                                300
glcb1_WT tccttggctg catcggttat ctgctgccac aacctgaaaa ggttaaggtc
glcb1_EG TCCTTGGCTG CATCGGTTAT CTGCTGCCAC AACCTGAAAA GGTTAAGGTC

           301                                                350
glcb1_WT ggcacggacc atgtcgaccg tgaggtagca gtgcaggccg gcccgcagtt
glcb1_EG GGCACGGACC ATGTCGACCG TGAGGTAGCA GTGCAGGCCG GCCCGCAGTT

           351                                                400
glcb1_WT ggtggtgccg gtgatgaacg cacgctatgc gctcaacgcc gctaatgccc
glcb1_EG GGTGGTGCCG GTGATGAACG CACGCTATGC GCTCAACGCC GCTAATGCCC

           401                                                450
glcb1_WT gttggggctc gttgtacgat gccctgtatg gcaccgacgc catcgccgaa
glcb1_EG GTTGGGGCTC GTTGTACGAT GCCCTGTATG GCACCGACGC CATCGCCGAA

           451                                                500
glcb1_WT gacggcggcg ccagcaaggg gcctggatac aatccggtgc gtggcgctcg
glcb1_EG GACGGCGGCG CCAGCAAGGG GCCTGGATAC AATCCGGTGC GTGGCGCTCG

           501                                                550
glcb1_WT agtcattgcg ttcgcccgcg ccttcctcga ccaggccttc cccctgcagg
glcb1_EG AGTCATTGCG TTCGCCCGCG CCTTCCTCGA CCAGGCCTTC CCCCTGCAGG

           551                                                600
glcb1_WT gcggttctca cgtcgatgcc agcgcctacc aggtagtgga tggtcaattg
glcb1_EG GCGGTTCTCA CGTCGATGCC AGCGCCTACC AGGTAGTGGA TGGTCAATTG

           601                                                650
glcb1_WT cgggtatcgc tgcatagcgg tgaagcgatc ggtttggccc aatcggatca
glcb1_EG CGGGTATCGC TGCATAGCGG TGAAGCGATC GGTTTGGCCC AATCGGATCA

           651                                                700
glcb1_WT attcgttggt caccagggcg aagcattgtc accgaccgtc gtactgctcg
glcb1_EG ATTCGTTGGT CACCAGGGCG AAGCATTGTC ACCGACCGTC GTACTGCTCG

           701                                                750
glcb1_WT tgcatcgtga cttgcacgtg gagatccaga tcgacccgac cagcccagtc
glcb1_EG TGCATCGTGA CTTGCACGTG GAGATCCAGA TCGACCCGAC CAGCCCAGTC

           751                                                800
glcb1_WT ggcagcaccg atgccgctgg cgtcaaggac ctggtgctgg agtcggcctt
```

```
glcb1_EG GGCAGCACCG ATGCCGCTGG CGTCAAGGAC CTGGTGCTGG AGTCGGCCTT

         801                                               850
glcb1_WT gaccaccatc atggactgcg aagattcagt ggccgcggtc gatgccgatg
glcb1_EG GACCACCATC ATGGACTGCG AAGATTCAGT GGCCGCGGTC GATGCCGATG

         851                                               900
glcb1_WT acaaggtggc tgtctaccgt aactggctgg gcctgatgaa gggcgacctg
glcb1_EG ACAAGGTGGC TGTCTACCGT AACTGGCTGG GCCTGATGAA GGGCGACCTG

         901                                               950
glcb1_WT actgaaaccg tgagcaagaa cgggaaggca ttcacgcgta ccctgaaccc
glcb1_EG ACTGAAACCG TGAGCAAGAA CGGGAAGGCA TTCACGCGTA CCCTGAACCC

         951                                              1000
glcb1_WT tgaccgcacc taccagggcc gcgacggtaa accagtaacc ctgccggggc
glcb1_EG TGACCGCACC TACCAGGGCC GCGACGGTAA ACCAGTAACC CTGCCGGGGC

         1001                                             1050
glcb1_WT gctcgctgct gttcgtgcgc aacgttggtc atctgatgag caccccaacc
glcb1_EG GCTCGCTGCT GTTCGTGCGC AACGTTGGTC ATCTGATGAG CACCCCAACC

         1051                                             1100
glcb1_WT atcctcgatc aggaaggtca ggagattccc gaaggaattc tcgatgccgt
glcb1_EG ATCCTCGATC AGGAAGGTCA GGAGATTCCC GAAGGAATTC TCGATGCCGT

         1101                                             1150
glcb1_WT aatcactagc cttgcagccg ttcacgacct gcgccgccgc ggcaactcgc
glcb1_EG AATCACTAGC CTTGCAGCCG TTCACGACCT GCGCCGCCGC GGCAACTCGC

         1151                                             1200
glcb1_WT gtagcggcag cgtctatatc gtgaagccca agatgcacgg ccctgctgag
glcb1_EG GTAGCGGCAG CGTCTATATC GTGAAGCCCA AGATGCACGG CCCTGCTGAG

         1201                                             1250
glcb1_WT gtagcatttg ccaacaggct gtttggtcga gttgaggatc ttcttggcct
glcb1_EG GTAGCATTTG CCAACAGGCT GTTTGGTCGA GTTGAGGATC TTCTTGGCCT

         1251                                             1300
glcb1_WT ggcccctcat accttgaaga tgggcatcat ggacgaggag cgccgcacca
glcb1_EG GGCCCCTCAT ACCTTGAAGA TGGGCATCAT GGACGAGGAG CGCCGCACCA

         1301                                             1350
glcb1_WT gcatcaacct caaggcgtgc atcgccgaag ccgccaaccg agtggtgttc
glcb1_EG GCATCAACCT CAAGGCGTGC ATCGCCGAAG CCGCCAACCG AGTGGTGTTC

         1351                                             1400
glcb1_WT atcaataccg ggttcctcga ccgtacgggc gacgagatgc actcgaccat
glcb1_EG ATCAATACCG GGTTCCTCGA CCGTACGGGC GACGAGATGC ACTCGACCAT

         1401                                             1450
glcb1_WT ggaagccggc gccatgctgc gcaagggcga catgaagtcg actccctgga
glcb1_EG GGAAGCCGGC GCCATGCTGC GCAAGGGCGA CATGAAGTCG ACTCCCTGGA

         1451                                             1500
glcb1_WT tccagtccta tgagcgcaac aacgtactgg tggggctgga ctgtggcctg
glcb1_EG TCCAGTCCTA TGAGCGCAAC AACGTACTGG TGGGGCTGGA CTGTGGCCTG

         1501                                             1550
```

```
glcb1_WT cgtggacggg cgcaaatcgg caagggcatg tgggccatgc cggacctgat
glcb1_EG CGTGGACGGG CGCAAATCGG CAAGGGCATG TGGGCCATGC CGGACCTGAT

         1551                                             1600
glcb1_WT ggcagccatg cttgaacaga aaattgctca ccccaaggcc ggcgccaaca
glcb1_EG GGCAGCCATG CTTGAACAGA AAATTGCTCA CCCCAAGGCC GGCGCCAACA

         1601                                             1650
glcb1_WT ctgcgtgggt accttcgcct accgccgcga cgctgcatgc gctgcattac
glcb1_EG CTGCGTGGGT ACCTTCGCCT ACCGCCGCGA CGCTGCATGC GCTGCATTAC

         1651                                             1700
glcb1_WT caccaagtag acgtgcgtca agtgcaacaa gagttggaac gcatcgacct
glcb1_EG CACCAAGTAG ACGTGCGTCA AGTGCAACAA GAGTTGGAAC GCATCGACCT

         1701                                             1750
glcb1_WT ggacagccag cgcgccgagt tgctgcaagg cctgctcagt gtgccggtca
glcb1_EG GGACAGCCAG CGCGCCGAGT TGCTGCAAGG CCTGCTCAGT GTGCCGGTCA

         1751                                             1800
glcb1_WT gtgtcgatcg caactggagc ccggccgaga tccaggcaga actggataac
glcb1_EG GTGTCGATCG CAACTGGAGC CCGGCCGAGA TCCAGGCAGA ACTGGATAAC

         1801                                             1850
glcb1_WT aactgccaga gcatccttgg ctatgtagtg cgctggatcg aacagggcgt
glcb1_EG AACTGCCAGA GCATCCTTGG CTATGTAGTG CGCTGGATCG AACAGGGCGT

         1851                                             1900
glcb1_WT tggttgttcc aaagtgctcg atatccacga cgtaggtttg atggaagacc
glcb1_EG TGGTTGTTCC AAAGTGCTCG ATATCCACGA CGTAGGTTTG ATGGAAGACC

         1901                                             1950
glcb1_WT gcgccacccct gcgtatctcc gcccagcata ttgcgaactg gctgcaccat
glcb1_EG GCGCCACCCT GCGTATCTCC GCCCAGCATA TTGCGAACTG GCTGCACCAT

         1951                                             2000
glcb1_WT cgtgtggtta gcactggcca agtccgtgca gcgctggagc gcatggctca
glcb1_EG CGTGTGGTTA GCACTGGCCA AGTCCGTGCA GCGCTGGAGC GCATGGCTCA

         2001                                             2050
glcb1_WT ggtggtcgat gggcagaatg ctggggacca agcctaccgc ccgatggcgc
glcb1_EG GGTGGTCGAT GGGCAGAATG CTGGGGACCA AGCCTACCGC CCGATGGCGC

         2051                                             2100
glcb1_WT cggactttga aagcagccat gccttccgcg ccgcctgtga cctggtgttc
glcb1_EG CGGACTTTGA AAGCAGCCAT GCCTTCCGCG CCGCCTGTGA CCTGGTGTTC

         2101                                             2150
glcb1_WT aagggacgtg agcagccaag tggctacaca gagccgctgc tccatgcctg
glcb1_EG AAGGGACGTG AGCAGCCAAG TGGCTACACA GAGCCGCTGC TCCATGCCTG

         2151                     2178
glcb1_WT gcggttacgc ttcaaacagg ctcgctga
glcb1_EG GCGGTTACGC TTCAAACAGG CTCGCTGA



         1                                                  50
glcb2_WT atgactggat acgttcaagt cggtggcctt caggtcgcca aggtcctgta
```

```
glcb2_EG ATAACTGAAT ACGTTTAAGT CGGTGGCCTT CAGGTCGCCA AGGTCCTGTA

         51                                               100
glcb2_WT cgacttcgtg aacaacgaag ccatccccgg gaccggcatc gtcgccgagc
glcb2_EG CGACTTCGTG AACAACGAAG CCATCCCCGG GACCGGCATC GTCGCCGAGC

         101                                              150
glcb2_WT agttctgggc gggtgcagag aagatcatca atgacctcgc tccaaagaac
glcb2_EG AGTTCTGGGC GGGTGCAGAG AAGATCATCA ATGACCTCGC TCCAAAGAAC

         151                                              200
glcb2_WT aaagccctgc tcgccaagcg cgacgagctg caagccaaga tcgacgcctg
glcb2_EG AAAGCCCTGC TCGCCAAGCG CGACGAGCTG CAAGCCAAGA TCGACGCCTG

         201                                              250
glcb2_WT gcaccaggca cgcaaaggcc aggcccacga cgccgcagcc tacaaagcat
glcb2_EG GCACCAGGCA CGCAAAGGCC AGGCCCACGA CGCCGCAGCC TACAAAGCAT

         251                                              300
glcb2_WT tcctccagga aatcggctac ctgctgccac aagccgacga tttccaggcc
glcb2_EG TCCTCCAGGA AATCGGCTAC CTGCTGCCAC AAGCCGACGA TTTCCAGGCC

         301                                              350
glcb2_WT accacccaga acgtggacga agaaatcgcc cacatggccg gcccacaact
glcb2_EG ACCACCCAGA ACGTGGACGA AGAAATCGCC CACATGGCCG GCCCACAACT

         351                                              400
glcb2_WT ggtcgttccg gtgatgaacg cccgcttcgc cctgaacgcc gccaacgccc
glcb2_EG GGTCGTTCCG GTGATGAACG CCCGCTTCGC CCTGAACGCC GCCAACGCCC

         401                                              450
glcb2_WT gctggggttc gctgtacgac gccctttacg gcaccgacgc catcagcgat
glcb2_EG GCTGGGGTTC GCTGTACGAC GCCCTTTACG GCACCGACGC CATCAGCGAT

         451                                              500
glcb2_WT gaaggcggcg ccgaaaaagg ccagggttac aacaaggtac gcggcgacaa
glcb2_EG GAAGGCGGCG CCGAAAAAGG CCAGGGTTAC AACAAGGTAC GCGGCGACAA

         501                                              550
glcb2_WT ggtcatcgca ttcgcccgcg ccttcctcga cgaagccgcg ccactggccg
glcb2_EG GGTCATCGCA TTCGCCCGCG CCTTCCTCGA CGAAGCCGCG CCACTGGCCG

         551                                              600
glcb2_WT ccggctcgca cgttgactcc acgggctacc gcatcgaagg cggcaagctg
glcb2_EG CCGGCTCGCA CGTTGACTCC ACGGGCTACC GCATCGAAGG CGGCAAGCTG

         601                                              650
glcb2_WT gttgttgcct tgaaaggcgg cagcaacagc ggcctgcgcg acgatgcgca
glcb2_EG GTTGTTGCCT TGAAAGGCGG CAGCAACAGC GGCCTGCGCG ACGATGCGCA

         651                                              700
glcb2_WT actgatcggc ttccacggcg acgccgccgc gccgactgcc gtgttgctca
glcb2_EG ACTGATCGGC TTCCACGGCG ACGCCGCCGC GCCGACTGCC GTGTTGCTCA

         701                                              750
glcb2_WT agcacaacgg cctgcacttc gaaatccagg tcgatgccag cacccccgtc
glcb2_EG AGCACAACGG CCTGCACTTC GAAATCCAGG TCGATGCCAG CACCCCCGTC

         751                                              800
```

```
glcb2_WT ggcagcaccg acgccgctgg cgtcaaagac atcctgatgg agtcggcact
glcb2_EG GGCAGCACCG ACGCCGCTGG CGTCAAAGAC ATCCTGATGG AGTCGGCACT

         801                                              850
glcb2_WT gaccaccatc atggactgcg aagactcggt tgcagccgta gacgctgacg
glcb2_EG GACCACCATC ATGGACTGCG AAGACTCGGT TGCAGCCGTA GACGCTGACG

         851                                              900
glcb2_WT acaaagtcat cgtctaccgc aactggctgg gcttgatgaa aggcgacctg
glcb2_EG ACAAAGTCAT CGTCTACCGC AACTGGCTGG GCTTGATGAA AGGCGACCTG

         901                                              950
glcb2_WT gccgaaagcg tgagcaaggg tggcaaaacc ttcacccgca ccatgaaccc
glcb2_EG GCCGAAAGCG TGAGCAAGGG TGGCAAAACC TTCACCCGCA CCATGAACCC

         951                                             1000
glcb2_WT ggaccgcgag tacgcggcgc ccaacggcgg cagcgtgacc ctgcacggtc
glcb2_EG GGACCGCGAG TACGCGGCGC CCAACGGCGG CAGCGTGACC CTGCACGGTC

         1001                                            1050
glcb2_WT gttcgttgct gttcgtgcgc aacgttggcc acctgatgac caacccggcg
glcb2_EG GTTCGTTGCT GTTCGTGCGC AACGTTGGCC ACCTGATGAC CAACCCGGCG

         1051                                            1100
glcb2_WT atcctcgatg cccaaggcaa cgaaatcccc gaaggtatcc aggacgggct
glcb2_EG ATCCTCGATG CCCAAGGCAA CGAAATCCCC GAAGGTATCC AGGACGGGCT

         1101                                            1150
glcb2_WT gttcaccaac ctgatcgccc tgcacaacct caacggcaac accagccgca
glcb2_EG GTTCACCAAC CTGATCGCCC TGCACAACCT CAACGGCAAC ACCAGCCGCA

         1151                                            1200
glcb2_WT agaatacccg cagcggcagc gtgtacatcg tcaagccgaa gatgcacggc
glcb2_EG AGAATACCCG CAGCGGCAGC GTGTACATCG TCAAGCCGAA GATGCACGGC

         1201                                            1250
glcb2_WT cctgaggaag tggccttcgc cgccgagatc ttcagccagg tcgaagacct
glcb2_EG CCTGAGGAAG TGGCCTTCGC CGCCGAGATC TTCAGCCAGG TCGAAGACCT

         1251                                            1300
glcb2_WT gctgggcatg ccgcgcaaca ccgtcaaggt cggcatcatg gacgaggaac
glcb2_EG GCTGGGCATG CCGCGCAACA CCGTCAAGGT CGGCATCATG GACGAGGAAC

         1301                                            1350
glcb2_WT gccgtaccac ggtcaacctc aaatcctgca tcaaggcagc cgccgagcgc
glcb2_EG GCCGTACCAC GGTCAACCTC AAATCCTGCA TCAAGGCAGC CGCCGAGCGC

         1351                                            1400
glcb2_WT gtggcgttca tcaacaccgg cttccttgac cgcactggcg atgaaatcca
glcb2_EG GTGGCGTTCA TCAACACCGG CTTCCTTGAC CGCACTGGCG ATGAAATCCA

         1401                                            1450
glcb2_WT cacctcgatg gaagccggcg ccgtggtgcg caaaggtgcc atgaagaacg
glcb2_EG CACCTCGATG GAAGCCGGCG CCGTGGTGCG CAAAGGTGCC ATGAAGAACG

         1451                                            1500
glcb2_WT agaagtggat cggcgcctac gaaaacaaca acgtcgacgt tggcctggcc
glcb2_EG AGAAGTGGAT CGGCGCCTAC GAAAACAACA ACGTCGACGT TGGCCTGGCC
```

```
          1501                                                1550
glcb2_WT accggcctgc aaggccgtgc gcagattggc aaaggcatgt gggccatgcc
glcb2_EG ACCGGCCTGC AAGGCCGTGC GCAGATTGGC AAAGGCATGT GGGCCATGCC

          1551                                                1600
glcb2_WT tgacctgatg gccgccatgc tcgagcagaa gatcgcccac ccgctggccg
glcb2_EG TGACCTGATG GCCGCCATGC TCGAGCAGAA GATCGCCCAC CCGCTGGCCG

          1601                                                1650
glcb2_WT gtgccaacac cgcttgggta ccgtcgccaa ctgccgccac cctgcacgcc
glcb2_EG GTGCCAACAC CGCTTGGGTA CCGTCGCCAA CTGCCGCCAC CCTGCACGCC

          1651                                                1700
glcb2_WT ctgcactacc acaaggtgga cgtacaggcg cgccagcgtg aactggcttc
glcb2_EG CTGCACTACC ACAAGGTGGA CGTACAGGCG CGCCAGCGTG AACTGGCTTC

          1701                                                1750
glcb2_WT acgtaccccg gcgtcggtgg acgacattct ggccattccg ctggctgccg
glcb2_EG ACGTACCCCG GCGTCGGTGG ACGACATTCT GGCCATTCCG CTGGCTGCCG

          1751                                                1800
glcb2_WT acaccaactg gtcggccgaa gagatccgca acgagctgga caacaacgcc
glcb2_EG ACACCAACTG GTCGGCCGAA GAGATCCGCA ACGAGCTGGA CAACAACGCC

          1801                                                1850
glcb2_WT cagggcattc tcggctacgt ggtgcgttgg atcgaccagg gcgtgggttg
glcb2_EG CAGGGCATTC TCGGCTACGT GGTGCGTTGG ATCGACCAGG GCGTGGGTTG

          1851                                                1900
glcb2_WT ctcgaaggtg ccggacatca acaacgtcgg cctgatggaa gaccgtgcca
glcb2_EG CTCGAAGGTG CCGGACATCA ACAACGTCGG CCTGATGGAA GACCGTGCCA

          1901                                                1950
glcb2_WT ccctgcgcat ctctgcccag ctgctggcca actggctgcg ccacggcgtg
glcb2_EG CCCTGCGCAT CTCTGCCCAG CTGCTGGCCA ACTGGCTGCG CCACGGCGTG

          1951                                                2000
glcb2_WT gtcagccagg agcaggtgct ggaaagcctc aagcgcatgg ccgtggtggt
glcb2_EG GTCAGCCAGG AGCAGGTGCT GGAAAGCCTC AAGCGCATGG CCGTGGTGGT

          2001                                                2050
glcb2_WT cgatcagcag aacgctggcg acgccctgta ccgcccgatg gcgccgaact
glcb2_EG CGATCAGCAG AACGCTGGCG ACGCCCTGTA CCGCCCGATG GCGCCGAACT

          2051                                                2100
glcb2_WT tcgacgacaa cgtggcgttc caggcggctg tggaactggt ggtggaaggt
glcb2_EG TCGACGACAA CGTGGCGTTC CAGGCGGCTG TGGAACTGGT GGTGGAAGGT

          2101                                                2150
glcb2_WT ggcaagcaac cgaacggtta taccgaaccg gtactgcacc gccgtcgccg
glcb2_EG GGCAAGCAAC CGAACGGTTA TACCGAACCG GTACTGCACC GCCGTCGCCG

          2151                         2178
glcb2_WT cgagttcaag gcgcgtaacg ggttgtaa
glcb2_EG CGAGTTCAAG GCGCGTAACG GGTTGTAA


        1                                                50
acea_WT atggcactga cacgcgaaca gcaaattgca gccctcgaga aagactgggc
```

```
acea_EG ATGGCACTGA CACGCGAACA GCAAATTGCA GCCCTCGAGA AAGACTGGGC

        51                                               100
acea_WT cgaaaacccg cgctggaaag gcgtgacccg tacctacacc gccgctgatg
acea_EG CGAAAACCCG CGCTGGAAAG GCGTGACCCG TACCTACACC GCCGCTGATG

        101                                              150
acea_WT tcgttcgcct gcgtggctcc gtgcaacctg agcacacctt tgcgcgtcag
acea_EG TCGTTCGCCT GCGTGGCTCC GTGCAACCTG AGCACACCTT TGCGCGTCAG

        151                                              200
acea_WT ggcgcagaaa aactgtggaa gctggtcacc gaaggtgccc acccgtcctt
acea_EG GGCGCAGAAA AACTGTGGAA GCTGGTCACC GAAGGTGCCC ACCCGTCCTT

        201                                              250
acea_WT ccgcccccgaa aaagatttcg tcaactgcat gggcgccctg acgggcggcc
acea_EG CCGCCCCGAA AAAGATTTCG TCAACTGCAT GGGCGCCCTG ACGGGCGGCC

        251                                              300
acea_WT aggctgtaca acaggtcaag gccggcatcc aggccatcta cctgtccggc
acea_EG AGGCTGTACA ACAGGTCAAG GCCGGCATCC AGGCCATCTA CCTGTCCGGC

        301                                              350
acea_WT tggcaggttg ccgccgacaa caactcggcc gagtcgatgt accctgacca
acea_EG TGGCAGGTTG CCGCCGACAA CAACTCGGCC GAGTCGATGT ACTTTGATTA

        351                                              400
acea_WT gtcgctgtac ccggtcgact cggtaccgac cgtggtcaaa cgcatcaaca
acea_EG GTCGCTGTAC CCGGTCGACT CGGTACCGAC CGTGGTCAAA CGCATCAACA

        401                                              450
acea_WT acgcgttccg ccgcgccgac cagatccagt ggaaagccgg caagaacccg
acea_EG ACGCGTTCCG CCGCGCCGAC CAGATCCAGT GGAAAGCCGG CAAGAACCCG

        451                                              500
acea_WT ggcgacgacg gctacatcga ctacttcgcg cccatcgtgg ccgacgccga
acea_EG GGCGACGACG GCTACATCGA CTACTTCGCG CCCATCGTGG CCGACGCCGA

        501                                              550
acea_WT agccggtttc ggtggcgtac tgaacgccta cgagctgatg aagaacatga
acea_EG AGCCGGTTTC GGTGGCGTAC TGAACGCCTA CGAGCTGATG AAGAACATGA

        551                                              600
acea_WT tcgaagcagg cgccgccggc gtgcacttcg aagaccagct ggcctcggtt
acea_EG TCGAAGCAGG CGCCGCCGGC GTGCACTTCG AAGACCAGCT GGCCTCGGTT

        601                                              650
acea_WT aaaaaatgcg gccacatggg cggcaaggtg ctggtaccga cccaggaagc
acea_EG AAAAAATGCG GCCACATGGG CGGCAAGGTG CTGGTACCGA CCCAGGAAGC

        651                                              700
acea_WT cgtacagaag ctggtagccg cgcgcctggc cgctgacgtg tcgggcgtac
acea_EG CGTACAGAAG CTGGTAGCCG CGCGCCTGGC CGCTGACGTG TCGGGCGTAC

        701                                              750
acea_WT cgaccatcat cctggcgcgt accgacgcca acgccgccga cctgctgacc
acea_EG CGACCATCAT CCTGGCGCGT ACCGACGCCA ACGCCGCCGA CCTGCTGACC

        751                                              800
```

```
acea_WT agcgactgcg acccgtacga ccagccgttc gtggttggcg agcgcacccg
acea_EG AGCGACTGCG ACCCGTACGA CCAGCCGTTC GTGGTTGGCG AGCGCACCCG

        801                                              850
acea_WT cgaaggcttc tacaaggtcc gtgccggcct cgatcaggcc attgcccgcg
acea_EG CGAAGGCTTC TACAAGGTCC GTGCCGGCCT CGATCAGGCC ATTGCCCGCG

        851                                              900
acea_WT gcctggccta cgccccgtac gccgacctga tctggtgtga aaccgccaag
acea_EG GCCTGGCCTA CGCCCCGTAC GCCGACCTGA TCTGGTGTGA AACCGCCAAG

        901                                              950
acea_WT ccggacctgg acgaagcccg ccgcttcgcc gaggcgatca agaaggagta
acea_EG CCGGACCTGG ACGAAGCCCG CCGCTTCGCC GAGGCGATCA AGAAGGAGTA

        951                                             1000
acea_WT cccggaccag atcctgtcgt acaactgctc gccttccttc aactggaaga
acea_EG CCCGGACCAG ATCCTGTCGT ACAACTGCTC GCCTTCCTTC AACTGGAAGA

        1001                                            1050
acea_WT aaaacctgga cgacgccacc atcgccaagt tccagcgcga attgtcggcc
acea_EG AAAACCTGGA CGACGCCACC ATCGCCAAGT TCCAGCGCGA ATTGTCGGCC

        1051                                            1100
acea_WT atgggttaca agcatcagtt catcaccctg gctggcatcc acaacatgtg
acea_EG ATGGGTTACA AGCATCAGTT CATCACCCTG GCTGGCATCC ACAACATGTG

        1101                                            1150
acea_WT gcacggcatg ttcaacctgg cgcacgacta cgcccgcaac gacatgaccg
acea_EG GCACGGCATG TTCAACCTGG CGCACGACTA CGCCCGCAAC GACATGACCG

        1151                                            1200
acea_WT cctacgtgaa gctgcaggag caggaattcg ctgacgccag caagggctac
acea_EG CCTACGTGAA GCTGCAGGAG CAGGAATTCG CTGACGCCAG CAAGGGCTAC

        1201                                            1250
acea_WT accttcgtgg cgcaccagca ggaagtgggc actggctact tcgacgacat
acea_EG ACCTTCGTGG CGCACCAGCA GGAAGTGGGC ACTGGCTACT TCGACGACAT

        1251                                            1300
acea_WT gaccaccgtg atccagggtg gcgcttcgtc ggtgactgca ctgaccggtt
acea_EG GACCACCGTG ATCCAGGGTG GCGCTTCGTC GGTGACTGCA CTGACCGGTT

        1301                     1326
acea_WT cgaccgagga agagcagttc cactga
acea_EG CGACCGAGGA AGAGCAGTTC CACTGA
```

**Figure S3.** Alignment of *P. putida* JM37 and *P. putida* JM37-ΔEG targeted knock-out genes for CRISPR edition. gRNA sequences are marked in yellow, target mutations in green and collateral mutations in red.

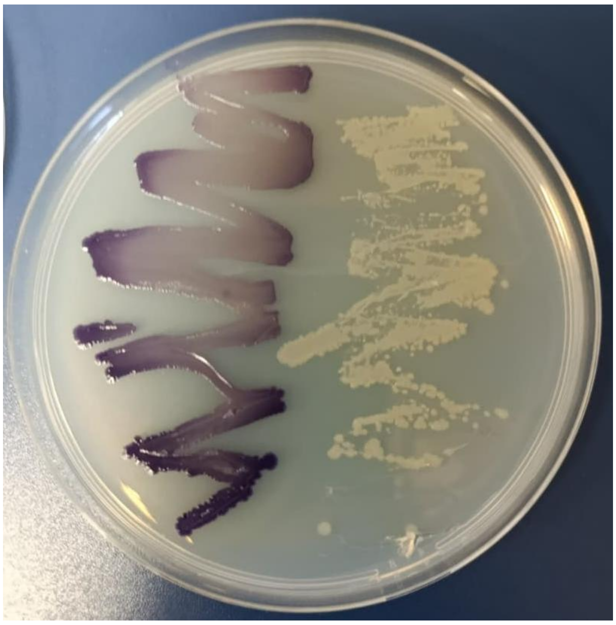

**Figure S4.** Crude violacein production by *P. putida* Vi strain in MC-Agar with 50 mM EG and 10 µM of N-(β-Ketocaproyl)-L-homoserine lactone as inducer (left). Wild-type strain is streaked as negative control (right).
